# Nuclear Myosin VI stabilises Ku-associated DNA ends during nonhomologous end joining

**DOI:** 10.64898/2026.09.01.748478

**Authors:** Abdulmohsen F. Alanazi, Alexander W. Cook, Isabel Shahid-Fuente, Natalia Fili, Tomas Venit, Ana-Maria Gherghelas, Noah B. Hitchcock, Alexandra Moreno Mitchell, Jesse Aaron, Teng-Leong Chew, Lin Wang, Piergiorgio Percipalle, Christopher P. Toseland

## Abstract

DNA double-strand breaks (DSBs) require rapid signalling and physical stabilisation of broken DNA ends to preserve genome integrity. Here, we identify myosin VI (MVI) as an ATM-regulated component of the DSB response. DNA damage induces rapid nuclear accumulation and nanoscale reorganisation of MVI across multiple cell models, in an ATM-dependent manner. Pharmacological or genetic perturbation of MVI attenuates γH2AX signalling and disrupts Ku80 organisation, while DNA damage persists. This leads to increased sensitivity to cisplatin and bleomycin. Super-resolution imaging reveals spatial association of MVI with Ku80-containing repair structures, implicating MVI in non-homologous end joining (NHEJ). In a minimal reconstituted system, MVI and actin enhance the proximity of Ku70/80-bound DNA ends. Together, our findings identify MVI as a regulator of DSB repair that links ATM signalling to Ku-associated DNA-end stabilisation and suggest that targeting MVI may sensitise tumour cells to genotoxic therapy.

## INTRODUCTION

DNA double-strand breaks (DSBs) are among the most cytotoxic forms of DNA damage and, if incorrectly repaired, they compromise genome stability (1). Cells therefore respond rapidly to DSB formation through coordinated signalling, chromatin remodelling and recruitment of specialised repair complexes. A central regulator of this response is ataxia-telangiectasia mutated (ATM) kinase, which is activated following DNA damage and phosphorylates multiple substrates involved in checkpoint signalling, chromatin organisation and DNA repair (2-4). One prominent consequence of ATM activation is phosphorylation of the histone variant H2AX at Ser139 to generate γH2AX, producing chromatin domains surrounding DNA lesions that facilitate recruitment and retention of DNA-damage-response proteins.

DSBs can be repaired through several pathways, principally homologous recombination and non-homologous end joining (NHEJ), with pathway utilisation influenced by cell-cycle stage and the molecular context of the lesion (2). NHEJ operates throughout much of the cell cycle and is initiated by the rapid recognition of exposed DNA ends by the Ku70– Ku80 heterodimer. Ku protects the broken ends and provides a platform for recruitment of DNA-PKcs and downstream repair factors including XRCC4 and DNA ligase IV (5). An important requirement during this process, is the maintenance of the two damaged DNA ends in sufficient proximity to permit productive synapsis and eventual ligation. Failure to constrain DNA-end movement can promote inappropriate repair and chromosome rearrangements. Consequently, DSB repair is not solely a biochemical process, but also represents a spatial and mechanical challenge in which chromatin mobility, nuclear architecture and repair-complex organisation must be carefully controlled.

Myosin VI (MVI) is an actin-dependent ATPase and depending upon its oligomeric state, binding partners and mechanical load, MVI can function both as a molecular transporter and as a force-sensitive anchor (6-8). In the cytoplasm, these properties support established functions in endocytosis, exocytosis and cell migration (9). However, a significant population of MVI also resides within the nucleus (10-13). Nuclear MVI associates with RNA polymerase II and transcriptional regulatory complexes and contributes to the spatial organisation and activity of transcription (8, 14). These mechanical properties raise the possibility that MVI could act as a general stabiliser of protein–DNA assemblies within the mechanically dynamic nuclear environment.

There are emerging indications that MVI may also participate in genome maintenance. Initial studies linked MVI to the p53 response following genotoxic stress, with p53 regulating MVI expression and MVI reciprocally influencing p53 stability and cell survival (15, 16). These effects appear to be highly context dependent, however, with contrasting responses reported between different cancer models (15, 16). More recently, MVI was shown to cooperate with WRNIP1 at stalled and reversed replication forks, where it protects nascent DNA from DNA2-dependent degradation during replication stress (17). Together, these observations suggest that nuclear MVI can contribute to genome protection. Whether MVI participates directly in canonical DSB repair, however, remains unresolved.

DNA damage is accompanied by changes in chromatin mobility, nuclear organisation and cellular mechanics, creating a requirement to maintain the spatial organisation of repair complexes while damaged chromatin is remodelled (18-22). A force-bearing motor, such as MVI, is ideally positioned to provide mechanical stabilisation during these large-scale changes and to support repair structures.

Defining such a role for MVI may also have implications for cancer therapy. MVI expression is elevated in several tumour types, including breast, ovarian and prostate cancers, and MVI has been associated with cancer-cell proliferation, migration and invasive behaviour (23-27). If MVI supports DNA repair, in addition to its established roles in tumour-associated transcription and signalling, disruption of MVI could therefore create an additional vulnerability to DNA-damaging therapies.

In this work, we identify MVI as an ATM-dependent component of the DSB response. DNA damage drives rapid nuclear accumulation and reorganisation of MVI. MVI associates with Ku80-containing repair structures, while its perturbation disrupts γH2AX signalling and Ku80 organisation, leading to persistent DNA damage and increased sensitivity to genotoxic treatment. We therefore propose that MVI functions as an ATM-regulated stabiliser of DNA ends, supporting their organisation during NHEJ and promoting efficient recovery from genotoxic stress.

## RESULTS

### DNA damage induces rapid ATM-dependent nuclear recruitment of myosin VI

To identify if MVI responds to DNA damage, immunofluorescence was used to determine the localisation of MVI before and after cisplatin or bleomycin treatment, across five mammalian cell lines representing tumorigenic (HeLa, MCF7, MCF10DCIS, MCF10CA1) and non-tumorigenic populations (MCF10a,). Cisplatin induces inter-strand and intra-strand cross-linking, which subsequently generates DSBs during DNA replication. Bleomycin however generates DSBs independent of the cell cycle stage.

In non-treated conditions, MVI is predominantly a cytoplasmic protein with a nuclear population (Figure 1 and Supplementary Figure 1). The level of nuclear MVI does vary across the cell lines with the highest found within MCF10DCIS cells. After cells have been treated with 65 µM cisplatin or 0.5 µM bleomycin for 4 hours, an increase in the the overall levels of MVI was observed, as previously reported (REF). Importantly, the MVI levels within the nucleus significantly increased across all cell lines (Figure 1C-D and Supplementary Figure 1B,D,F). Measurement of the nuclear:cytoplasmic (N:C) ratio ensured that the observed enrichment is due to the nuclear recruitment of protein, rather than a change in total protein levels. The presence of DNA damage was confirmed by staining for γH2AX and foci determination (Figure 1A-D and Supplementary Figure 1).

**Figure 1.**
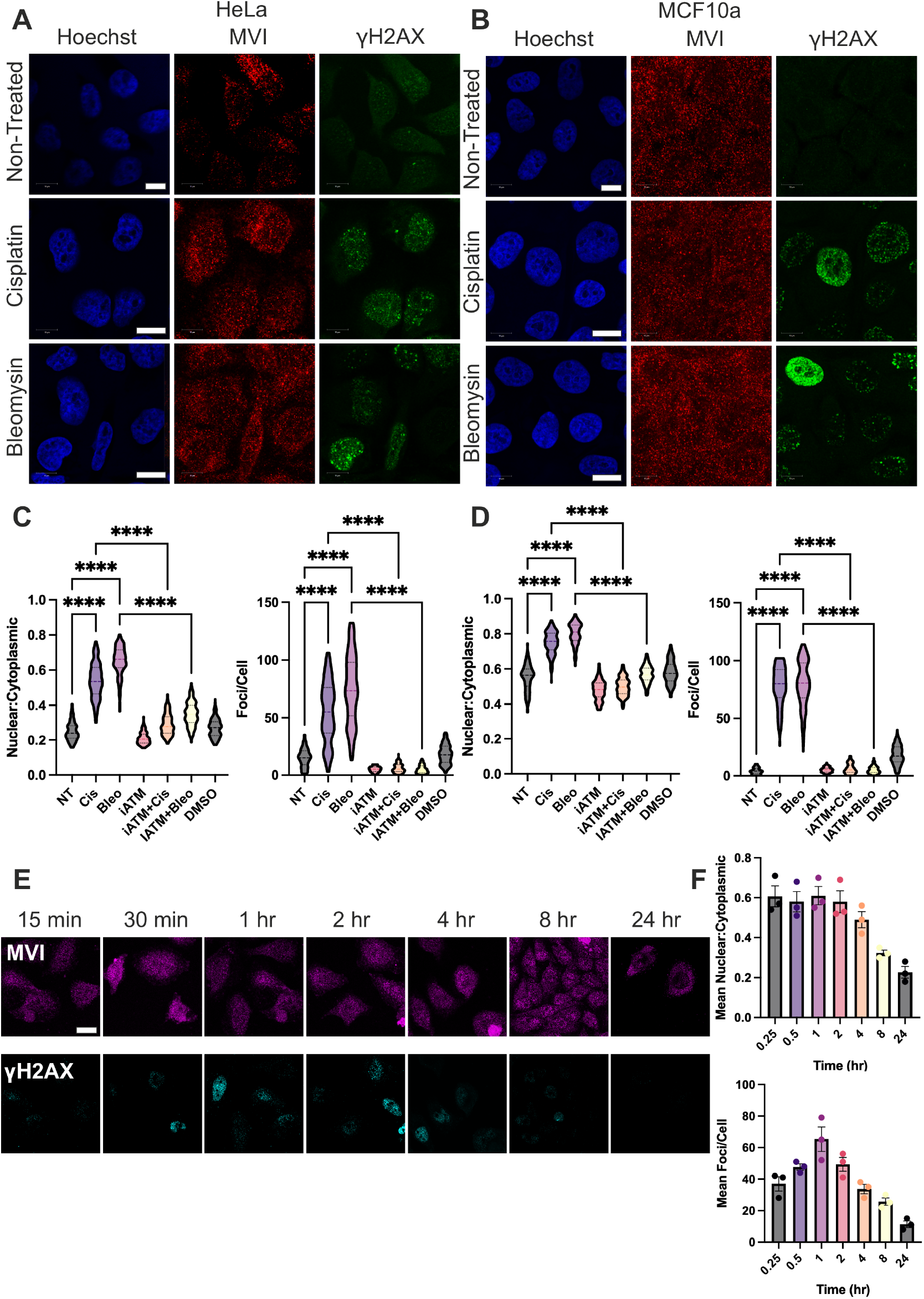
The nuclear accumulation of myosin VI following DNA damage. Representative Immunofluorescence staining against MVI (red), γH2AX (green) and DNA (blue) in **(A)** HeLa cells and **(B)** MCF10a cells for non-treated, cisplatin (65 µM 4 h) and bleomycin (0.5 µM 4 hr) treated cells (Scale bar 10 μm). **(C)** and **(D)** Quantification of nuclear MVI and number of γH2AX foci for HeLa and MCF10a cells, respectively. 150 cells per condition from three independent experiments. ****p <0.0001 by one-way ANOVA and Tukey’s multiple comparisons test. Representative images for the iATM (20 µM for 4 h) and DMSO treatments are in Supplementary Figure 2. **(E)** Representative Immunofluorescence staining against MVI (magenta) and γH2AX (cyan) at the indicated timepoints following exposure to 65 µM cisplatin (Scale bar 10 μm). **(F)** Quantification of nuclear MVI and number of γH2AX foci for the time course measurement. Each datapoint represents the mean from 40 cells. Errors bars represent SEM.

The use of ATM kinase inhibitor (Ku55933) in HeLa and MCF10a cells blocked the recruitment of MVI and as expected, depleted the γH2AX signal in the presence of DNA damage (Figure 1C-D and Supplementary Figure 2). This suggests that the nuclear recruitment of MVI is tied to the DNA damage response pathway.

To understand the timings of nuclear recruitment, a time-course measurement was performed in HeLa cells. The cells were treated with cisplatin, as above, but a selection of cells were fixed at intervals between 15 min and 24 h (Figure 1E). The samples were then stained for MVI and γH2AX before nuclear levels and foci counting was performed (Figure 1F). The level of MVI immediately rises to a mean N:C of 0.6 from a baseline of 0.25. The number of γH2AX foci rise for the first hour then subsides to baseline after 24 h. The rapid kinetics of MVI recruitment suggest that it participates at an early stage of the DNA-damage response.

### MVI role in the DNA damage response is transcription-independent

As MVI has a well-established role in RNA polymerase II transcription, we next asked whether MVI affects the DNA damage response through transcriptional regulation of relevant genes. To address this, we performed bulk RNA-seq analysis of WT and MVI knockdown HeLa cells treated with cisplatin (25 µM for 24 h) and compared them with the corresponding untreated controls (Supplementary Figure 3).

First, comparison of MVI knockdown cells with WT cells identified 1947 differentially expressed genes, comprising 1458 downregulated and 489 upregulated genes (Supplementary Figure 3B), consistent with previous findings (8). However, gene ontology analysis of these differentially expressed genes did not identify significantly enriched terms associated with the DNA damage response or DNA repair (Supplementary Figure 3B). This suggests that MVI role in these processes is independent of its role in DNA transcription

Cisplatin treatment of WT HeLa cells induced a broad transcriptional response, with 1816 genes downregulated and 2291 genes upregulated (Supplementary Figure 3D). Many of these genes were associated with the DNA damage response, as indicated by gene ontology analysis (Supplementary Figure 3E). Cisplatin treatment of MVI knockdown cells resulted in a largely similar transcriptional response, with only 210 genes differentially expressed compared with cisplatin-treated WT cells. These genes were associated with GO terms related to cell growth and proliferation, but not with the DSB response (Supplementary Figure 3F and 3G). These findings further support the conclusion that the role of MVI in the DNA damage response is largely independent of its role in transcription.

### DNA damage reorganises nuclear myosin VI into large damage-associated structures

As previously reported, MVI forms distinct clusters within the cell nucleus with a proportion associated with RNA Polymerase II (8). To understand the role of MVI in the DNA damage response, we performed Stochastic Optical Reconstruction Microscopy (STORM) and cluster analysis to measure the spatial organisation of MVI in HeLa cell nuclei. Following the treatment of cells with cisplatin (65 µM 4 h), we observed changes in the nanoscale organisation of nuclear MVI (Figure 2A).

**Figure 2.**
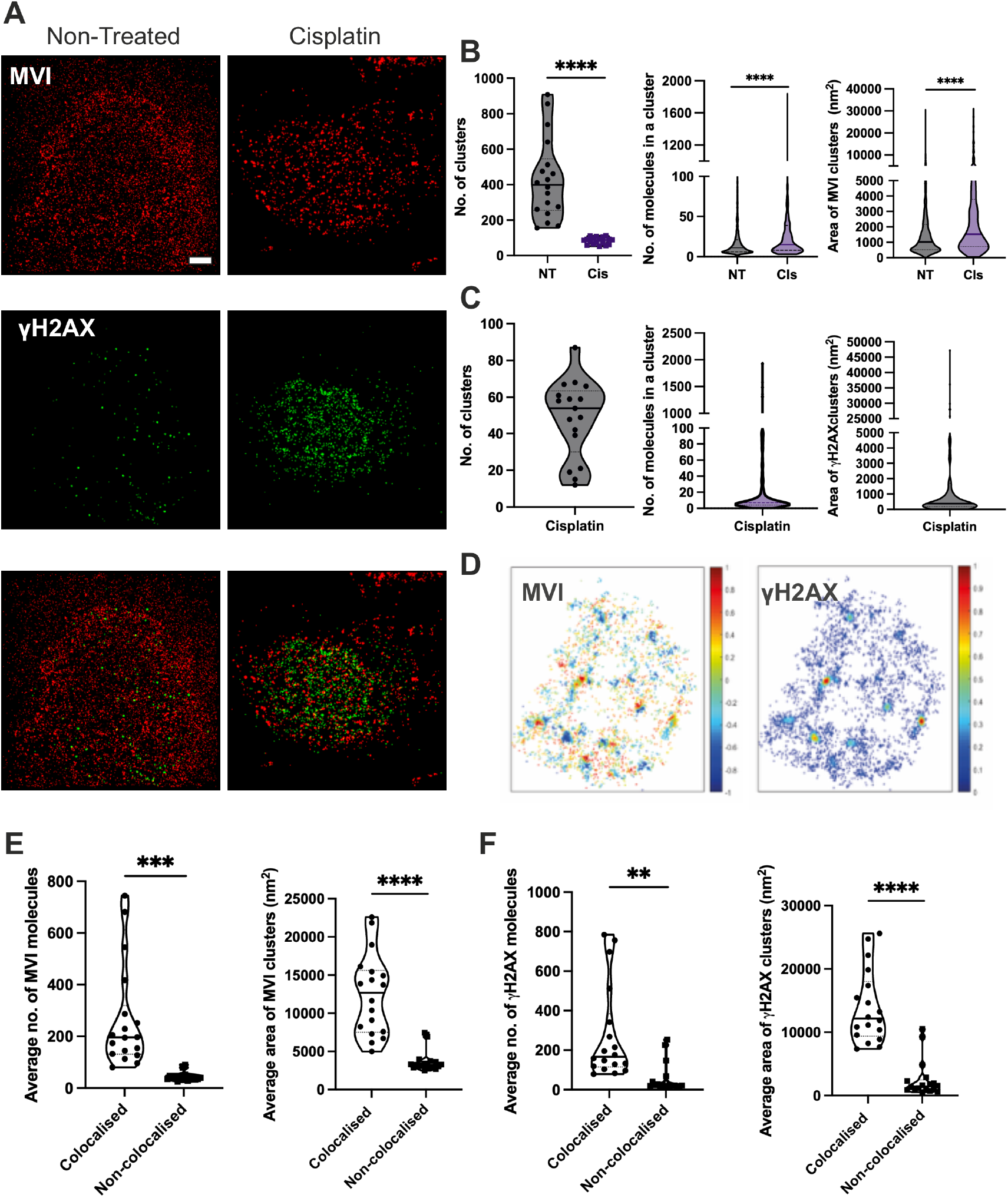
Myosin VI is associated with large γH2AX structures. **(A)** Example STORM render images of MVI and γH2AX under non-treated and cisplatin treatment (65 µM 4 h) in HeLa cells (scale bar 2 μm). The γH2AX staining denotes the nucleus region which was taken forward for cluster analysis. **(B)** Cluster analysis of MVI nuclear organisation. Individual data points correspond to the average value for a cell ROI (n = 17). ****p <0.0001 by one-way ANOVA and Tukey’s multiple comparisons test. **(C)** Cluster analysis of γH2AX nuclear organisation. Individual data points correspond to the average value for a cell ROI (n = 17). **(D)** Representative colocalisation heatmap of MVI and γH2AX clusters following cisplatin treatment/ DoC values of 1 are perfectly colocalised and -1 are separated from each other. **(E)** Colocalisation cluster analysis for MVI. **(F)** Colocalisation cluster analysis for γH2AX. The depicted by black circles values represent the mean from the ROIs for each protein. **p <0.01 ***p <0.001 ****p <0.0001 by one-way ANOVA and Tukey’s multiple comparisons test. Error bars represent SEM.

In untreated cells, an average of 400 MVI clusters per cell were identified with an average of 23 molecules within each cluster, and an average cluster size of 1025 nm^2^ (Figure 2B). However, after treatment with cisplatin, the number of clusters within a nucleus decreased to 85. Interestingly, the number of MVI molecules per cluster and cluster area significantly increased. This highlights that MVI is responding to DNA damage, possibly altering its role from transcription to DNA damage. As expected, cisplatin treatment induced increased molecular clustering of γH2AX (Figure 2A, C).

To determine if this organisational change of MVI coincides with the location of the DSB signal staining for γH2AX, cluster colocalization between the two proteins was measured using the Degree of Colocalisation (DoC) analysis (28). This analysis employs a coordinate-based method to determine correlation between individual molecules in each channel (29). Overall, there is no systematic colocalization between the two proteins. However, further interrogation of the data revealed that, whilst the small, dispersed clusters do not colocalise, the larger clusters do display colocalisation (represented as signal above 0) (Figure 2D). Quantification showed that the average number of MVI and γH2AX molecules in a cluster, and their cluster area were significantly higher in colocalised structures, compared to the non-colocalised ones (Figure 2E-F).

Taken together, we conclude that both proteins colocalise at large damage sites which are typically associated with the repair location (30-32). Moreover, the extent of the colocalization is likely to vary during the time course following damage until repair is completed. Here, our analysis was performed at the 4 h timepoint, to ensure that the potential repair sites are captured. However, as shown in Figure 1E, after 4 h treatment, the number of γH2AX foci has already decreased, as repair is underway. Therefore, the observed co-localisation might be enhanced at earlier timepoints.

### ATM signalling regulates MVI mobilisation and nuclear recruitment following DNA damage

While STORM provided a static snapshot of MVI, we examined the dynamic behaviour of MVI in live HeLa cells in the absence and presence of cisplatin treatment. We used the aberration-corrected multi-focal microscope (acMFM) system (33) to simultaneously track single-molecules across nine focal planes in live-cells (Figure 3A), covering 4 μm in the *z* axis and 20 × 20 μm in *xy*. In this way, we were able to observe and track the 3D dynamics of stably expressed halo-tagged MVI in the cytoplasm and nucleus. 3D trajectories displayed a clear increase in movement within the cytoplasm following treatment with cisplatin (Figure 3B). To quantify our observations, we determined the diffusion constant for each track by measuring the Mean Squared Displacement (MSD) and then plotted the average diffusion constant per cell for the cytoplasm and nucleus (Figure C-D). Cisplatin and Bleomycin treatment caused a significant increase in mean cytoplasmic MVI diffusion with an increase from 0.3 µm^2^/s to 0.7 µm^2^/s. Whilst the nuclear diffusion was also significantly increased, the change was minor compared to the cytoplasm 0.2 µm^2^/s to 0.3 µm^2^/s. This nuclear diffusion is consistent with the STORM imaging data, where the nucleus has less MVI clusters following DNA damage, suggesting that more molecules are in a freely diffusing state. Moreover, this is consistent with our previous measurements showing cisplatin treatment causing an overall increase in diffusion (19).

**Figure 3.**
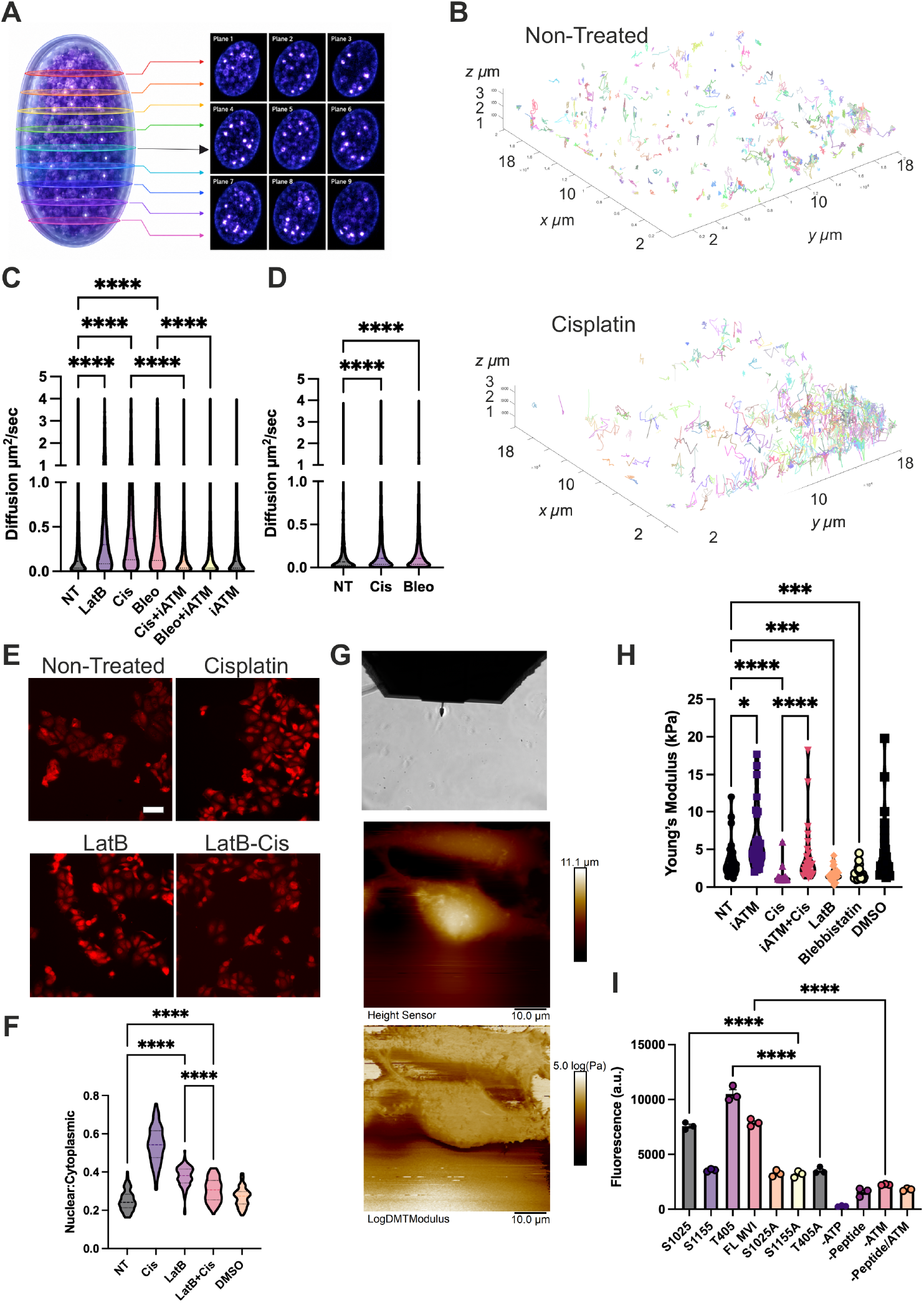
ATM kinase regulates Myosin VI nuclear localisation. **(A)** Illustration depicting simultaneous acquisition of 9 focal planes covering 4 μm in the z-axis to perform live cell 3D single molecule tracking of fluorescently tagged proteins. **(B)** Representative example render of 3D single molecule trajectories for non-treated and cisplatin treated HeLa cells. **(C)** and **(D)** Plot of Halo-MVI diffusion constants under the stated conditions derived from fitting trajectories to an anomalous diffusion model, as described in methods for cytoplasm and nucleus, respectively. Combined data from 50 ROIs across three independent experiments. ****p <0.0001 by one-way ANOVA and Tukey’s multiple comparisons test. **(E)** Representative Immunofluorescence staining against MVI (red) under the stated conditions. (Scale bar 40 μm) **(F)** Quantification of nuclear recruitment of MVI, showing the N:C ratio under the stated conditions. Each datapoint represents the mean from 100 cells. ****p <0.0001 by one-way ANOVA and Tukey’s multiple comparisons test. **(G)** Representative brightfield image of HeLa cells with the AFM probe, a topography render depicting cell height and shape, and a render of nanomechanical map across the cell. **(H)** Mean Young’s modulus values for the stated conditions where each point represents a single cell (n=25 from three independent experiments). *p <0.05 ***p <0.001 ****p <0.0001 by one-way ANOVA and Tukey’s multiple comparisons test. **(I)** Quantification of ATM kinase activity using substrate peptides and full length (FL) MVI. Experiments were performed in triplicate. The -ATP, -ATM and -peptide/ATM represents background signal within the reporter. -Peptide reflects background signal within the reporter and non-specific ATM kinase activity. ****p <0.0001 by one-way ANOVA and Tukey’s multiple comparisons test.

Focusing on the cytoplasmic population of MVI, we propose that the rapid diffusion facilitates the nuclear relocation. We have previously identified that relatively static MVI is functional, so this diffusive species would not be expected to be performing a function within the cytoplasm (34). This is further confirmed through the use of an ATM inhibitor which blocked the DNA damage response and MVI nuclear recruitment. Under these conditions, MVI cytoplasmic diffusion did not increase, consistent with the diffusion increase being linked to nuclear localisation.

Interestingly, the level of diffusion increase caused by the DNA damage agents is similar to cytoskeletal and mechanical perturbation via treatment with Latrunculin B (LatB). We subsequently explored if LatB treatment directly leads to nuclear localisation of MVI (Figure 3E-F). Indeed, there was a significant increase in nuclear MVI following treatment, but at a lower level compared to cisplatin treatment. This suggests that the DNA damage recruitment utilises several signalling and/or targeting approaches. Of note, treatment with a combination of LatB and cisplatin perturbed the nuclear recruitment of MVI. Mechanistically, this observation may indicate either the need of actin filaments to support the nuclear localisation of MVI or that the combined drug treatments cause a stress response which halts the nuclear recruitment.

We next tested whether ATM-dependent MVI recruitment was associated with changes in cellular mechanics. Atomic Force Microscopy (AFM) was used to map stiffness across HeLa cells in non-treated and treated conditions (Figure 3G-H). Consistent with previous reports, cisplatin caused a decrease in stiffness which was perturbed by use of the ATM kinase inhibitor, consistent with previous findings (19). As expected, LatB and blebbistatin also caused a decrease in stiffness. Interestingly, the use of the ATM inhibitor alone caused an increase in cell stiffness, as reported previously (35). These observations link MVI localisation to changes in cellular stiffness, with softer cells having levels of nuclear MVI.

Nevertheless, as suggested by Figure 3F and H, a decrease in stiffness alone (LatB treatment) does not yield the same magnitude of MVI nuclear recruitment as cisplatin treatment. We hypothesised that MVI nuclear recruitment is under the control of a signalling event. Therefore, we explored if the ATM kinase can directly phosphorylate MVI. MVI has several potential phosphorylation sites, but there is limited information about their functional significance. Here, we used an *in vitro* kinase activity assay using recombinant ATM, along with peptides for potential phosphorylation sites and full length MVI (Figure 3I). Control peptides, where the potential phosphorylation targets were mutated to alanine, were used to establish background activity. S1025 and T405, along with full length MVI, where substrates for the ATM kinase, highlighting that the kinase can phosphorylate MVI. These data identify MVI as a potential direct ATM substrate and suggest that ATM may regulate MVI through a combination of direct phosphorylation and changes in cellular mechanics.

### Perturbation of Myosin VI disrupts the DNA damage response

We have observed a rapid recruitment of MVI and its localisation to large γH2AX foci (Figure 1). We next turned our attention to the functional consequences of MVI within the DNA damage response. To this end, we have performed siRNA knockdown of MVI in HeLa and MCF10a cells. Interestingly, following MVI knockdown, there was a lack of γH2AX response following cisplatin treatment (Figure 4).

**Figure 4.**
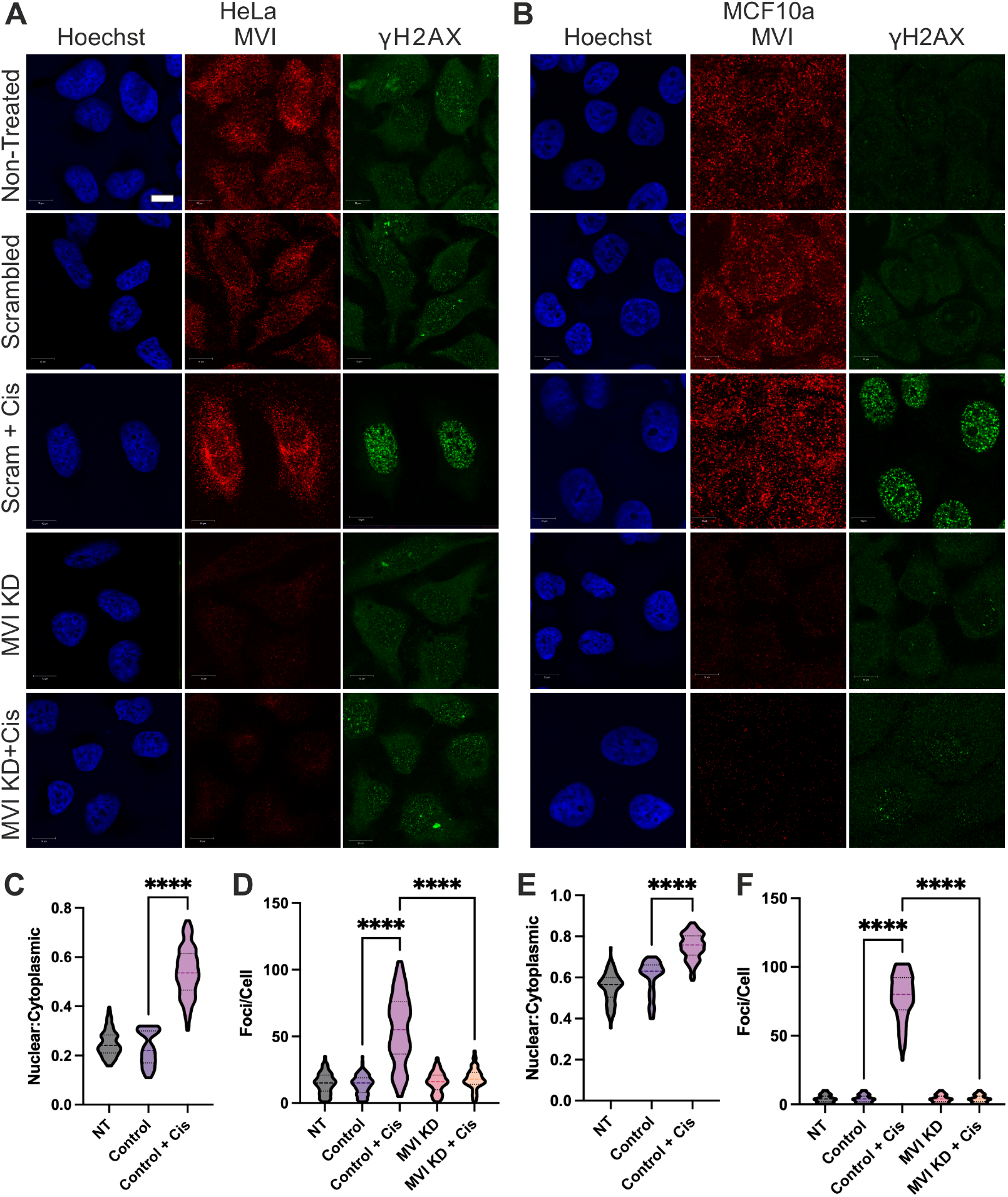
Myosin VI knockdown disrupts γH2AX signalling. Representative Immunofluorescence staining against MVI (red), γH2AX (green) and DNA (blue) in **(A)** HeLa cells and **(B)** MCF10a cells for non-treated and cisplatin (65 µM 4 h) treated cells (Scale bar 10 μm). The cisplatin treatment was conducted in the presence of MVI siRNA treatment (MVI KD) and control siRNA treatment (Scrambled). **(C)** and **(D)** Quantification of nuclear MVI (except for siRNA treatment) and number of γH2AX foci for HeLa and MCF10a cells, respectively. 150 cells per condition from three independent experiments. ****p <0.0001 by one-way ANOVA and Tukey’s multiple comparisons test.

To further explore this effect, we used a small molecule inhibitor of MVI, TIP (2,4,6-triiodophenol), which has been shown to disrupt its transcriptional roles (8, 11, 14, 36, 37). TIP was used in combination with the DNA damage agents, cisplatin and bleomycin in HeLa and MCF10a cells, and cisplatin alone in MCF7, MCF10CA1 and MCF10DCIS (Figure 5 and Supplementary Figure 5).

**Figure 5.**
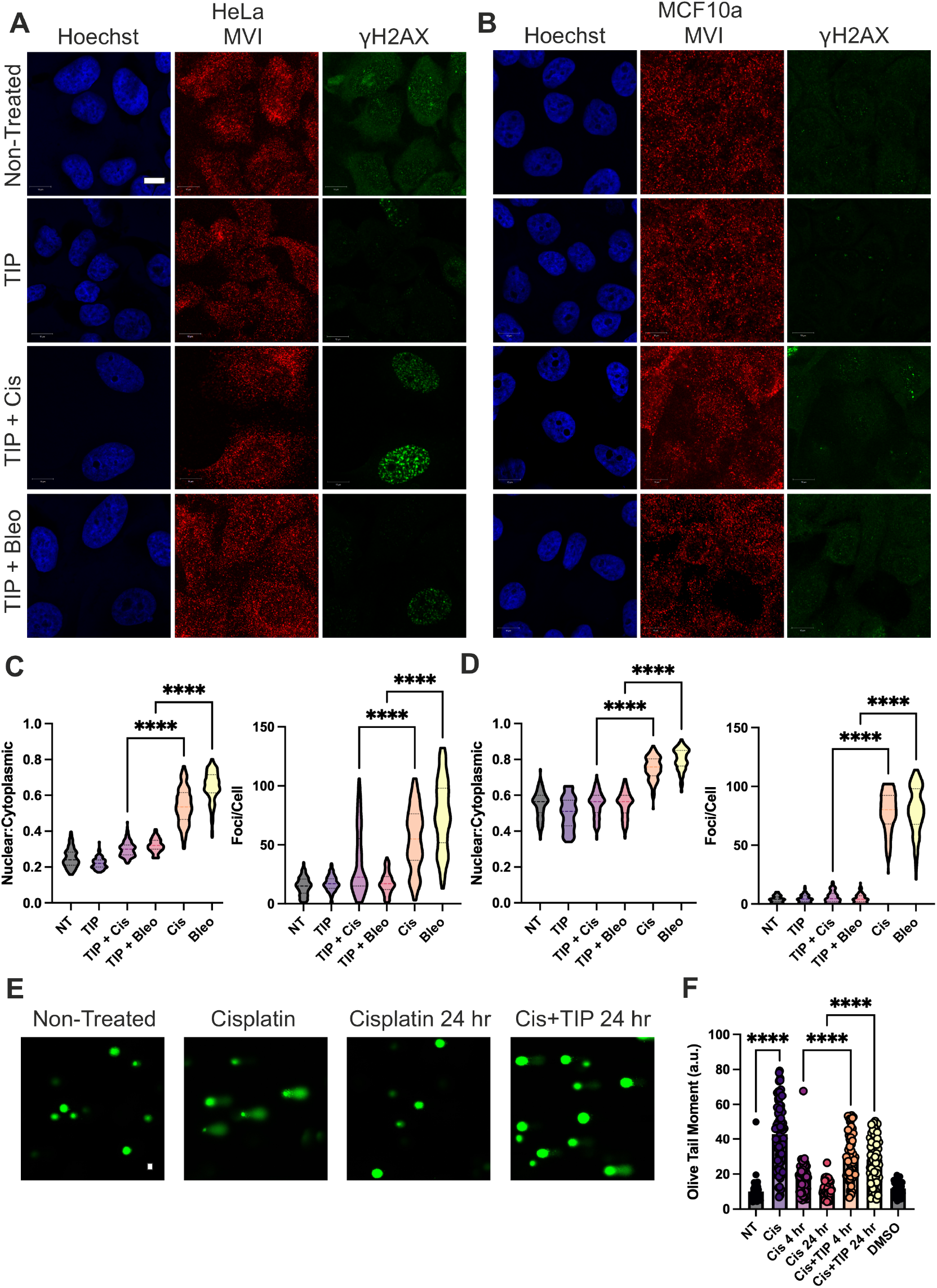
Myosin VI perturbation disrupts γH2AX signalling and inhibits DNA repair. Representative Immunofluorescence staining against MVI (red), γH2AX (green) and DNA (blue) in **(A)** HeLa cells and **(B)** MCF10a cells for non-treated, TIP-treated (25 µM 4 h), TIP+cisplatin (65 µM 4 h) and TIP+bleomycin (0.5 µM 4 hr) treated cells (Scale bar 10 μm). **(C)** and **(D)** Quantification of nuclear MVI and number of γH2AX foci for HeLa and MCF10a cells, respectively. 150 cells per condition from three independent experiments. ****p <0.0001 by one-way ANOVA and Tukey’s multiple comparisons test. (E) Representative alkaline comet assay images, DNA stained with Vista green dye. (Scale bar = 20 µm). **(F)** Quantification of alkaline comet assay analysis measuring olive tail moments in the stated conditions (At least 50 cells per condition). ****p <0.0001 by one-way ANOVA and Tukey’s multiple comparisons test.

With respect to MVI nuclear localisation (Figure 5C, E and Supplementary Figure 5C, D, F), TIP suppressed the recruitment (MCF10a, MCF10CA1 and MCF10DCIS) or reduced it, in the case of HeLa and MCF7. Nevertheless, all cell lines displayed less recruitment compared to cisplatin treatment alone. Across all conditions, we found that TIP treatment perturbed the γH2AX responses (Figure 5 C, D, and Supplementary Figure 5C, D, F). However, the extent of the response did vary across the cell line population, which likely reflects diversity in the cell lines and potential differential activity of TIP in those cell lines.

As both siRNA knockdown and TIP impact the whole cellular pool of MVI, we specifically targeted the nuclear pool of MVI by over-expression of a dominant negative MVI NLS-Cargo binding domain in HeLa cells (Supplementary Figure 6A and B). The construct is strongly expressed in the nucleus and sequesters protein interactors thereby displacing MVI. Consistent with the knockdown and TIP treatment, perturbation of the nuclear pool of MVI attenuated the γH2AX response following cisplatin treatment.

Previous work has established that MVI perturbation can impact histone marks associated with transcription whereby there is an increase in heterochromatin when MVI is inhibited (8). Here, we have stained HeLa cells for three heterochromatin marks (H3K9me3, H3K27me3 and H4K20me1) in the presence and absence of TIP (Supplementary Figure 6C). In all cases, the levels of heterochromatin increased following TIP treatment (Supplementary Figure 6D). Interestingly, heterochromatin suppresses γH2AX (38), raising the possibility that altered chromatin state contributes to the attenuated γH2AX response following MVI inhibition.

Lastly, COMET assays were employed to directly monitor the presence of DNA damage to ensure that damage still occurs despite the perturbed γH2AX signalling (Figure 5E). Following Cisplatin treatment in the presence of TIP, DSBs are still generated, as seen by the increase in tail moment (Figure 5F). Intriguingly, the damage remains unrepaired at 24 h in the presence of TIP, suggesting that the lack of functional MVI causes a significant repair defect.

### Myosin VI inhibition sensitises cells to genotoxic treatment

The COMET assays suggest that TIP co-treatment leads to prolonged DNA damage which has the potential to increase cell death. To explore the relationship between MVI perturbation and cell death, we performed cell viability (MTT) assays and screened Live-Dead cells through microscopy following 24 h treatment.

All cell lines were treated with cisplatin and bleomycin, in the absence and presence of TIP. Cytotoxicity increased in the presence of TIP and was significantly increased during the combination of Cis+TIP, over monotherapy for all cell lines except MCF7 (Figure 6A). Nevertheless, even in MCF7, there was a non-significant increase in cytotoxicity. A following titration experiment was performed in HeLa cells treated with varying concentrations of cisplatin and a fixed 25 µM TIP (Figure 6B). The cisplatin IC_50_ value decreased from 59 µM to 10 µM in the presence of TIP.

**Figure 6.**
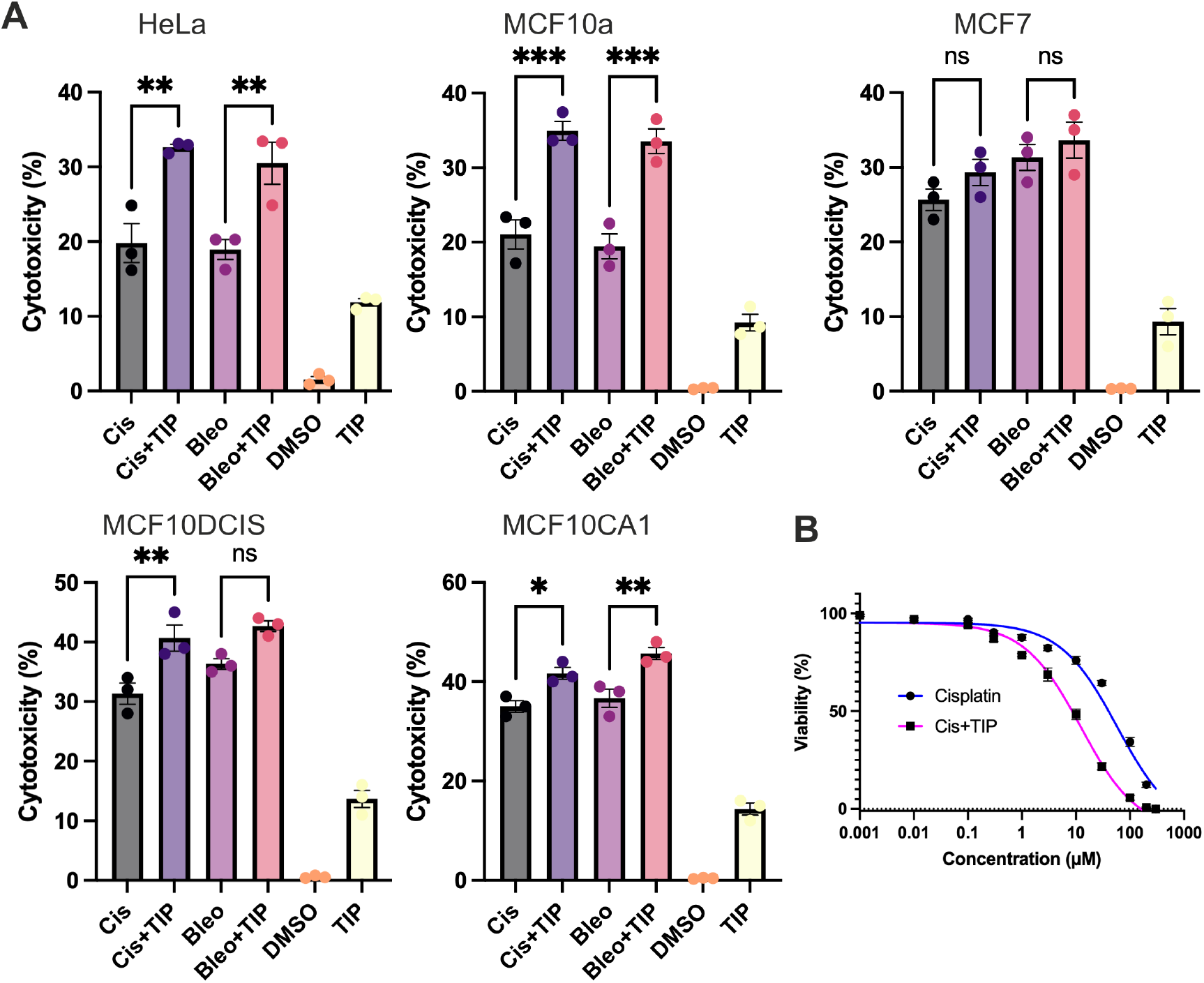
Myosin VI inhibition sensitises cells to DNA damage agents. **(A)** MTT cell viability assay under the stated conditions for each cell line. Treatments were performed for 24 hrs. The datapoints represent the mean from five technical repeats across three biological repeats. All data are normalised to non-treated control. *p <0.05, **p <0.01, ***p <0.001 by one-way ANOVA and Tukey’s multiple comparisons test. (B) MTT assay titration of cisplatin in the presence and absence of 25 µM TIP. IC50 fitting yields 59 µM and 10 µM in the absence and presence of TIP, respectively. Error bars represent SEM.

A Live-Dead screen was employed to directly measure cell death and then extended the cell lines to A375 (melanoma), SHEP21N (neuroblastoma), LN18 (glioblastoma) and T24 (bladder carcinoma). HeLa and MCF7 were included for comparison to the MTT assay (Supplementary Figure 7). Consistent with the MTT assay, combined treatment of HeLa cells with Cisplatin and TIP resulted in a significant increase in cell death, whilst MCF7 cells did not show a significant increase. Interestingly, all the additional cell lines tested also displayed an increase in cell death following the combined treatment. The largest effect was observed for LN18 glioblastoma and T24 bladder carcinoma models, where cell death increased more than four-fold.

### Myosin VI stabilises Ku-associated DNA ends during NHEJ

Whilst we have reported that MVI perturbation can disrupt γH2AX signalling and increase cell death, the function of MVI within the DNA damage response pathway has not been defined.

Given the rapid response of MVI to the DNA damage, we focused on how MVI supports DNA repair, within the NHEJ pathway. Moreover, previous proteomic analysis linked MVI to this pathway through the interaction with Ku80 (39). Ku80 forms a heterodimer with Ku70 to bind the free DNA ends within a DSB. It is critical for the complex to stabilise the free DNA ends to prevent genomic rearrangements and the generation of chromosome fusions.

Using STORM, we measured the clustering behaviour of MVI and Ku80 (Figure 7A). Following DNA damage there was an increase in the number and size of Ku80 clusters (Figure 7B). However, these foci were disrupted when MVI was perturbed via TIP, with the Ku80 clusters still being present, but in smaller numbers. Morevoer, significant colocalization was observed between MVI and Ku80 following cisplatin treatment as evidenced by the heatmap (Figure 7C) and histogram of Degree of Colocalisation (DoC) scores, whereby Ku80 colocalised with MVI and MVI colocalised with Ku80 (Figure 7D).

**Figure 7.**
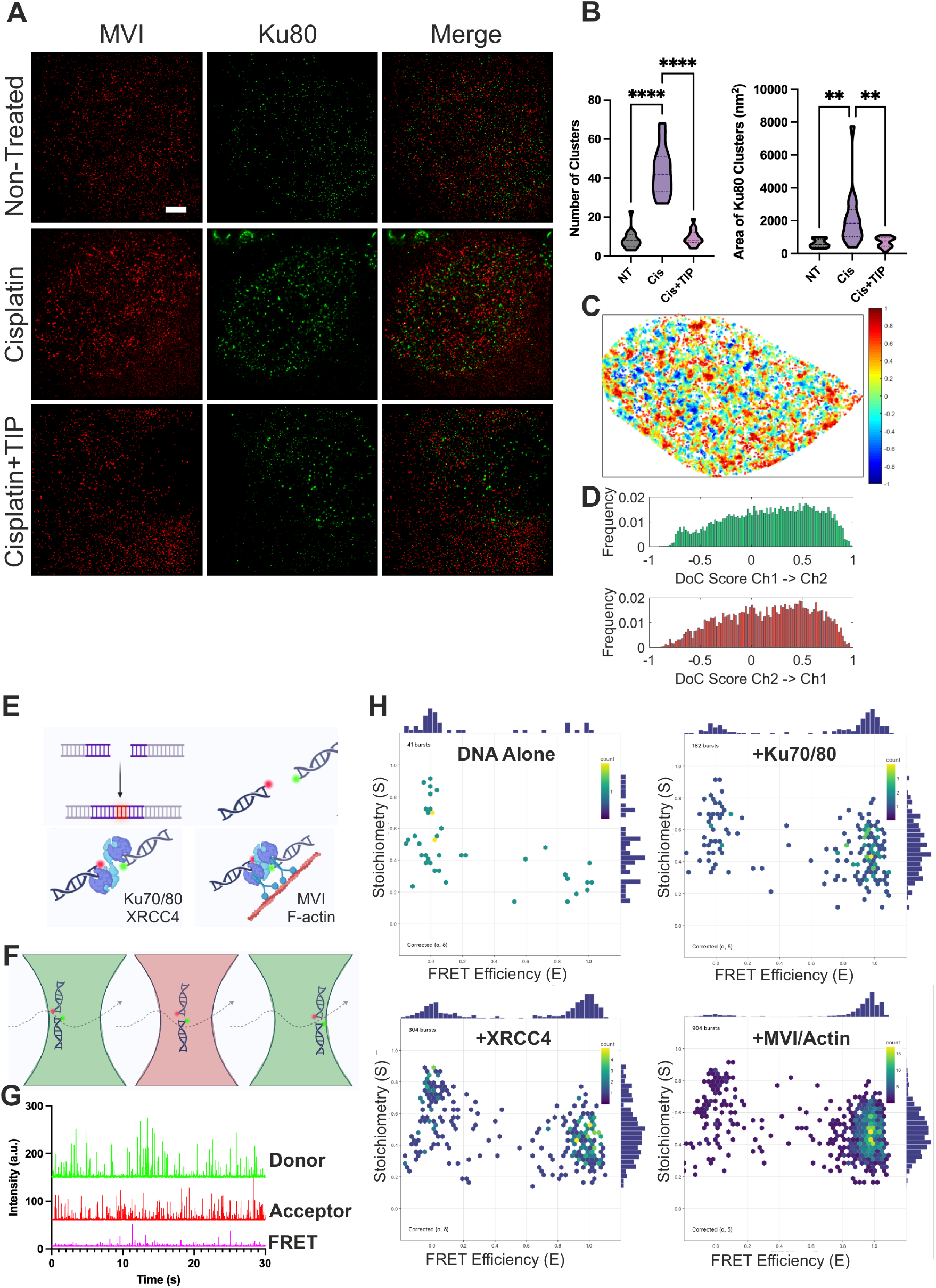
Myosin VI stabilises Ku80 at the synaptic complex to support DNA repair. **(A)** Example STORM render images of MVI and Ku80 under non-treated, cisplatin treatment (65 µM 4 h) and combined Cisplatin+TIP (65 µM and 25 µM 4 h) in HeLa cells (scale bar 2 μm). The Ku80 staining denotes the nucleus region which was taken forward for cluster analysis. **(B)** Cluster analysis of Ku80 nuclear organisation. Individual data points correspond to the average value for a cell ROI (n = 17). ****p <0.0001 by one-way ANOVA and Tukey’s multiple comparisons test. **(C)** Representative colocalisation heatmap of MVI and Ku80 clusters following cisplatin treatment/ DoC values of 1 are perfectly colocalised and -1 are separated from each other. **(D)** Histogram of DoC scores for Ku80 colocalised with MVI and MVI colocalised with Ku80. **(E)** Illustration of the DNA oligonucleotide, labelled with Atto550 and AlexaFluor647, designed for an *in vitro* DSB. The proteins are presented in hypothetical arrangements to bridge the DNA oligonucleotides to generate a FRET signal. **(F)** Representation of ALEX smFRET, whereby when the two oligonucleotides diffuse into the confocal volume, the lasers alternate the excitation during the transit, allowing the determination of smFRET and the presence of both dyes. **(G)** Representation of photon bursts for Donor detector, Acceptor detector and the resulting FRET burst. **(H)** Plots of FRET efficiency vs stoichiometry for 20 pM DNA oligonucleotides in the presence of 500 nM Ku70/80, then 500 nM XRCC4 **(I)** and 500 nM MVI plus 1 µM F-actin **(J). (K)** The number of FRET bursts for each experimental condition stated. **p<0.01 ****p <0.0001 by one-way ANOVA and Tukey’s multiple comparisons test. Error bars represent SEM.

This implies that MVI may support the localisation of Ku80 at damage-associated assemblies. To investigate this relationship further, we turned to an *in vitro* minimalised system using recombinant proteins consisting of MVI, filamentous actin (F-actin), Ku70/80 and XRCC4 (scaffold protein), similar to previous studies (5). We used fluorescently labelled double-stranded oligonucleotides to mimic a DSB. Proteins were hypothesised to bind to this duplex creating a bridge resulting in a high FRET signal (Figure 7E). We monitored the joining of the DNA duplexes using confocal-based alternating laser excitation single molecule FRET (smFRET). In this way, as the oligonucleotides pass through the confocal volume, we can measure FRET and simultaneously ensure both labels are present by recording their photon bursts (Figure 7F-G).

The addition of Ku70/80 caused a partial increase in the high FRET species suggesting bridging of the DNA duplexes (Figure 7H), which was further enhanced by the addition of XRCC4 (Figure 7I). However, the presence of MVI and F-actin caused a significant increase in the number of high FRET events (Figure 7J). The number of events were quantified (Figure 7K), whereby we observe that MVI and actin are both required to enhance the proximity between the DNA ends. Moreover, the use of the MVI spring mutant (8), which cannot undergo force-induced anchoring, failed to induce the association between the DNA duplexes, thereby highlighting that MVI can promote the close juxtaposition of Ku-bound DNA ends in an anchoring-related manner.

## DISCUSSION

Here, we identify MVI as an ATM-regulated component of the cellular response to DNA damage and provide evidence that it supports repair through the organisation and stabilisation of Ku70/80-associated DNA ends (Figure 8). Following DNA damage, MVI rapidly accumulated within the nucleus across multiple cell line models. Nuclear recruitment was accompanied by altered molecular mobility, reorganisation of MVI clusters and selective association with large γH2AX-positive structures. Perturbation of MVI impaired γH2AX signalling, disrupted Ku80 organisation, prolonged the persistence of DNA damage and increased sensitivity to genotoxic treatment. Collectively, these findings support a model in which MVI acts as a regulator of DSB repair, promoting the stable juxtaposition of broken DNA ends during NHEJ.

**Figure 8.**
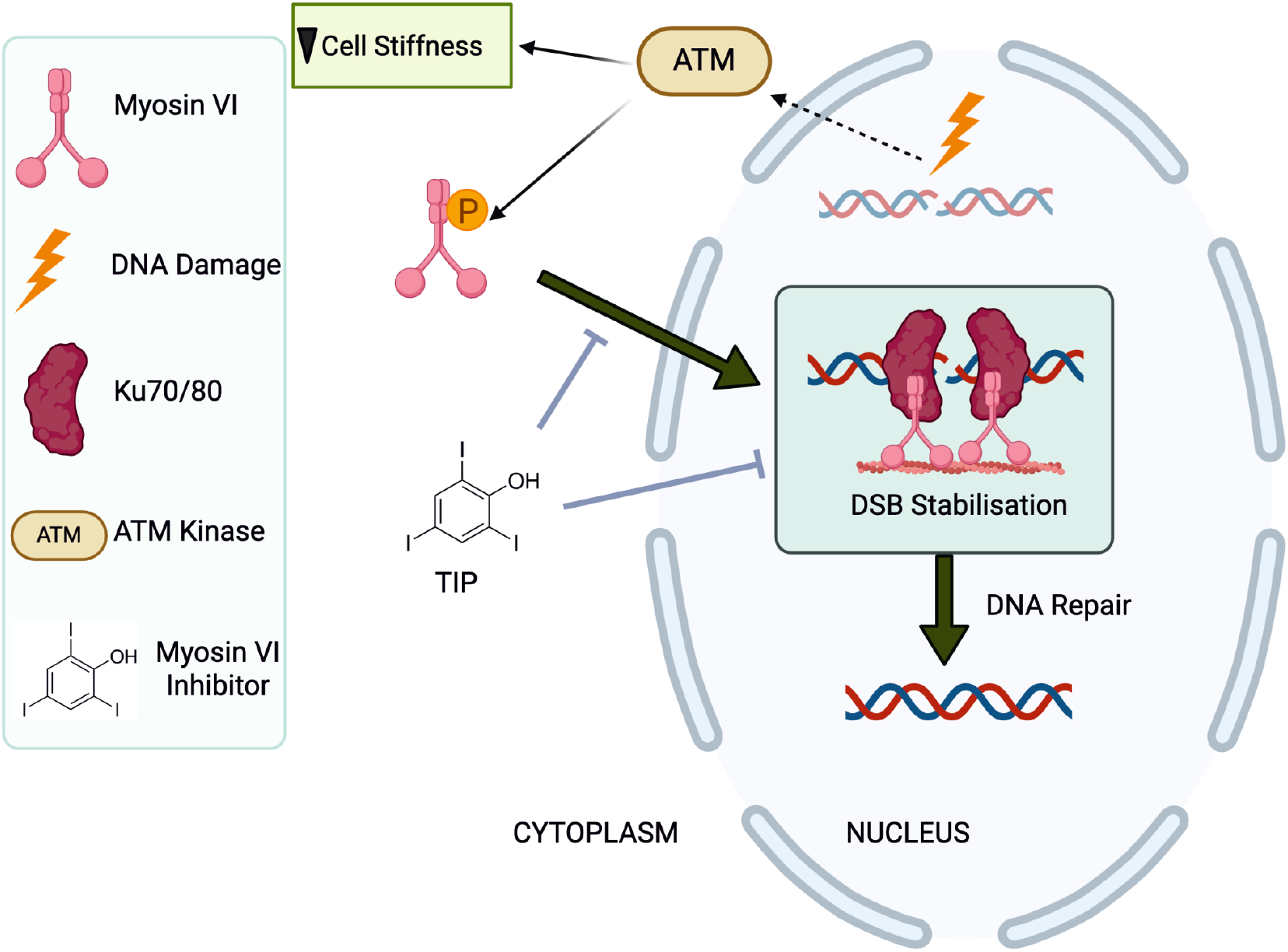
Schematic of myosin VI stabilisation of Double-strand breaks. Based on the data presented here we can propose the following model: DNA damage activates the ATM kinase which phosphorylates MVI and triggers a decrease in cell stiffness. This causes the rapid nuclear recruitment of MVI to DSB sites and interactions with Ku80. This stabilisation of Ku80 facilitates DNA repair via NHEJ. TIP inhibitions of MVI disrupts the nuclear localisation of MVI and its stabilisation of Ku80. The nucleus undergoes significant mechanical and chromatin remodelling during the DNA damage response. Failure to stabilise the DSB causes the ends to lose contact thereby prolonging the DSB and eventually causing cell death. The figure was generated in Biorender.

The rapid nuclear accumulation of MVI suggests that it participates relatively early in the DNA-damage response. This occurred independent of the damage agent, despite the different mechanisms through which they generate DNA lesions. ATM appears to regulate this response. Pharmacological inhibition of ATM prevented the DNA-damage-induced increase in nuclear MVI and suppressed the associated increase in cytoplasmic MVI mobility. The *in vitro* kinase experiments further demonstrated that MVI can serve as ATM substrate. Phosphorylation could alter MVI conformation, cargo-binding activity or interaction with actin, thereby promoting release from cytoplasmic complexes and subsequent nuclear entry. In addition to direct biochemical regulation, the results indicate that ATM-dependent changes in cellular mechanics may contribute to MVI mobilisation. Cisplatin treatment reduced cellular stiffness, increased cytoplasmic MVI diffusion and promoted nuclear accumulation, whereas ATM inhibition attenuated each of these responses. Disruption of actin polymerisation with Latrunculin B similarly reduced stiffness and induced a more modest increase in nuclear MVI. This suggests that mechanical softening or cytoskeletal remodelling is sufficient to mobilise part of the MVI population but does not reproduce the complete DNA-damage response. A plausible model is therefore that MVI recruitment requires both damage-specific signalling and a permissive mechanical state.

Super-resolution imaging showed that DNA damage not only increased nuclear MVI levels but also substantially changed its nanoscale organisation. Untreated nuclei contained numerous relatively small MVI clusters, consistent with the previously described involvement of MVI in transcriptional organisation (8). Following cisplatin treatment, cluster number decreased while cluster area and the number of molecules per cluster increased. This redistribution suggests that MVI is reorganised from multiple basal nuclear assemblies into fewer, larger structures associated with the damage response. Such behaviour may represent a functional transition from transcription-associated complexes towards repair-associated assemblies. This is further supported by RNA-seq evidence showing that MVI depletion does not impact damaged-induced gene expression changes.

The heterogeneous relationship between MVI and γH2AX may reflect selective recruitment of MVI to a subset of lesions, including complex, persistent or actively repairing breaks. It is also important to recognise that γH2AX domains extend across chromatin surrounding a lesion rather than precisely marking the physical DNA end. Consequently, a repair factor acting directly at or close to the break would not necessarily display uniform colocalisation with the entire γH2AX domain.

The evidence for a direct function within DNA repair comes from the relationship between MVI and Ku80. Ku70 and Ku80 form the DNA-end-binding heterodimer that initiates classical NHEJ and protects broken ends from excessive processing or inappropriate rearrangement (5). Following DNA damage, MVI colocalised with Ku80, and MVI perturbation reduced the number and altered the organisation of Ku80 clusters. These findings suggest that MVI contributes to the assembly, retention or stability of Ku70/80 repair complexes. SmFRET experiments support a model whereby MVI could stabilise Ku70/80 at DNA ends, promote higher-order organisation of Ku-bound lesions or provide a physical linkage between the repair machinery and an actin-based nuclear scaffold.

This proposed function is consistent with how MVI can function as a force-sensitive anchor when mechanical load is applied across the motor. At a DNA break, MVI may therefore act as a molecular tether. By coupling Ku70/80 DNA ends to actin, MVI could resist separation of the two broken ends and maintain their spatial alignment while repair proceeds. Such a mechanism would be particularly valuable within the mechanically dynamic nuclear environment, where chromatin motion could otherwise increase the risk of inappropriate end joining, chromosome translocations or loss of genetic material. Although the results strongly implicate MVI in NHEJ, it remains possible that MVI also influences other repair processes or acts at lesions that can be processed through more than one pathway.

Perturbation of MVI also attenuated the γH2AX response despite the continued presence of DNA damage. The COMET assays showed that lesions were still generated and persisted for at least 24 hours when MVI was inhibited. MVI inhibition also increased H3K9me3, H3K27me3 and H4K20me1, suggesting broader changes in chromatin state. Increased chromatin compaction could limit access of ATM to H2AX, restrict propagation of γH2AX domains or interfere with recruitment of repair factors. Therefore, the impact upon γH2AX could be indirect through broader chromatin changes.

The persistence of DNA damage following MVI perturbation was accompanied by increased DNA damage sensitivity. MVI has established functions in membrane trafficking, cell migration and transcription. Therefore, MVI inhibition could affect normal tissues, as well as tumours. However, MVI is elevated in several tumours providing a targeting advantage (23-27). In ER-positive breast cancer, MVI inhibition may offer an additional therapeutic opportunity because MVI contributes to both

ER-dependent transcription (14) and survival following DNA damage. Simultaneous disruption of oncogenic transcriptional activity and DNA repair could provide a mechanistically attractive approach.

Together, these findings identify MVI as an ATM-regulated component of the DSB response that promotes the organisation and stable proximity of Ku-bound DNA ends. This expands the nuclear functions of MVI beyond transcription and replication-fork protection and identifies a mechanochemical contribution to DNA-end synapsis during NHEJ.

## ACKNOWLEDGEMENTS

We thank the UKRI-MRC (MR/M020606/1) UKRI-BBSRC (BB/X008460/1), UKRI-STFC (19130001) and the Royal Society (IES\R3\183138 and SIF\R2\252076) for funding to C.P.T. Aberration-corrected multi-focal microscopy was performed in collaboration with the Advanced Imaging Center at Janelia Research Campus, a facility supported by the Howard Hughes Medical Institute. We also thank Satya Khuon (Janelia Research Campus) for assisting with cell culture. The JF549 dyes were kindly provided by Luke Lavis (Janelia Research Campus). We also thank grant from NYU Abu Dhabi to P.P. and we acknowledge technical help from the NYU Abu Dhabi Center for Genomics and Systems Biology, in particular Marc Arnoux and Mehar Sultana. We appreciate the computational platform provided by NYUAD HPC team and are especially thankful to Nizar Drou for technical help.

## AUTHOR CONTRIBUTIONS

C.P.T. conceived the study. A.F.A, A.W.C., I.S-F, N.F., T.V., A-M.G., N.B.H, A.M.M. and C.P.T. designed and performed experiments. N.F. and C.P.T. designed and cloned constructs. A.W.C., N.F. and C.P.T. performed single molecule imaging experiments. Imaging was supported by L.W. and J.A. and T-L.C. A.W.C., T.V., and P.P. performed and analyzed the genomics experiments. N.F. and C.P.T. expressed, purified and performed experiments with recombinant proteins. C.P.T. supervised the study. C.P.T. wrote the manuscript with comments from all authors.

## MATERIALS AND METHODS

### Constructs

pcDNA3.1 Halo-MVI, pFastBac MVI, pFastBac MVI Spring and pcDNA3.1 Halo-NLS-CBD was generated previously (40).

### Protein Expression using Baculovirus system

Full-length MVI Non-insert, MVI Spring and *Xenopus* calmodulin were expressed in *Sf9* and *Sf21* (*Spodoptera frugiperda*) insect cells using the Bac-to-Bac® Baculovirus Expression System (Invitrogen). *Sf9* cells were cultured in Sf900 media (Gibco). Recombinant bacmids were generated following the manufacturer’s instructions and were transfected into adherent Sf9 cells to generate the P1 viral stock. *Sf9* cells were infected in suspension at 27ºC and 100 rpm with 1 in 50 dilution of P1 and P2 viral stocks to yield P2 and P3 stocks, respectively. Finally, expression of recombinant proteins was set up by infecting *sf21* cells with the P3 viral stock in Spodopan media (PAN Biotech). To ensure correct folding of the myosin VI constructs, cells were simultaneously infected with P3 viral stock of the myosin VI constructs together with calmodulin at a 0.75 ratio The cells were harvested after 3 days by centrifugation for 15 min at 700xg and at 4 °C and resuspended in ice cold myosin extraction buffer (90 mM KH_2_PO_4_, 60 mM K_2_HPO_4_, 300 mM KCl, pH 6.8), supplemented with Proteoloc protease inhibitor cocktail (Expedeon) and 100 µM PMSF, before proceeding to protein purification. Prior to sonication, an additional 5 mg recombinant calmodulin was added together with 2 mM DTT. After sonication, 5 mM ATP and 10 mM MgCl_2_ were added and the solution was rotated at 4 °C for 30 min before centrifugation (20,000g, 4°C, 30 min). Then, the cell lysate was subjected to the purification. Proteins were purified by affinity chromatography (HisTrap FF, GE Healthcare). The purest fractions were further purified through a Superdex 200 16/600 column (GE Healthcare).

### Cell culture and Transfection

HeLa (ECACC 93021013), MCF7 (ECACC 86012803) cells were cultured at 37ºC and 5% CO_2_, in Gibco MEM Alpha medium with GlutaMAX (no nucleosides), supplemented with 10% Fetal Bovine Serum (Gibco), 100 units/ml penicillin and 100 µg/ml streptomycin (Gibco). For the transient expression of NLS-CBD, HeLa cells grown on glass coverslips were transfected using Lipofectamine 2000 (Invitrogen), following the manufacturer’s instructions. Depending on the construct, 48 h after transfection, cells were subjected to drug treatment then nuclear staining using Hoechst 33342 (Thermo Scientific), fixed and analysed or subjected to indirect immunofluorescence (see below). MCF10 cells (ATCC CRL-10317) were maintained in 50% MEM Alpha media with GlutaMAX (no nucleosides) and 50% Ham’s F-12 nutrient mixture, supplemented with 5% FBS, 5% horse serum, penicillin-streptomycin mix diluted to 100 units mL^-1^, 50µg Cholera toxin, 5µg insulin, 20ngmL^-1^ human EGF and 0.5µgmL^-1^ hydrocortisone. MCF10DCIS (a gift from Dr Elena Rainero) cells were cultured in DMEM/Ham’s F-12 (1:1) medium, supplemented with 5% horse serum, 10 µg/mL insulin, 20 ng/mL EGF (Sigma, Cat. No. E9644), 0.5 mg/mL hydrocortisone, and 1% PS. MCF10CA1 cells (a gift from Dr Elena Rainero) were cultured using the same formulation, with hydrocortisone reduced to 0.2 mg/mL. A375, SHEP21N, LN18 and T24 cells were a gift from Dr Greg Wells and cultured in MEM, see HeLa above.

### Cell Treatments

For MVI knock-down experiments, HeLa cell monolayers, seeded to 30 – 50 % confluency, were transfected with human myosin VI siRNA duplex (5′GGUUUAGGUGUUAAUGAAGtt-3′) (Ambion) or AllStars Negative Control siRNA duplex (Qiagen) at a concentration of 50 nM, using Lipofectamine 2000 (Invitrogen), according to the manufacturer’s guidelines. Cells were fixed or harvested after 48 h for further analysis. To inhibit myosin VI, cells were treated with 25 µM TIP (Sigma) for 1 h at 37 ºC. To inhibit actin polymerization, cells were treated with 1 µM Latrunculin B (Sigma) for 1 h. Bleomycin sulphate (Sigma, Cat no.B1141000) was prepared to a stock concentration of 100µM is dH_2_0 and used at a concentration of 0.5 µM for 4 h. Cisplatin (Sigma, Cat no. C2210000) was prepared in 0.9% NaCl solution to a stock concentration of 1mg.mL^-1^ and used at a concentration of 65 µM for 4 h unless stated. ATM inhibitor Ku55933 (Sigma, Cat no. SML1109) was prepared in DMSO, to a stock concentration of 5mg.mL^-1^ and used at 20 µM for 4 h. Blebbistatin (Sigma) was resuspended to 50 mM in DMSO and used at a concentration of 50 µM for 1– 2 h.

### Immunofluorescence

Cells were seeded onto 13-mm sterile circular glass (N1.5) coverslips at a density of 160k cell/ml and cultured as above for 24-hours to allow cell attachment. Cells were then subjected to the stated treatments. Cells were fixed for 15 min at room temperature in 4% (w/v) paraformaldehyde (PFA) in PBS and residual PFA was quenched for 15 min with 50 mM ammonium chloride in PBS. All subsequent steps were performed at room temperature. Cells were permeabilised and simultaneously blocked for 15 min with 0.1 % (v/v) Triton X-100 and 2 % (w/v) BSA in PBS. Cells were then immuno-stained against the endogenous proteins by 1 h incubation with the indicated primary and subsequently the appropriate fluorophore-conjugated secondary antibody (details below), both diluted in 2 % (w/v) BSA in PBS.

The following antibodies were used at the indicated dilutions: rabbit anti-myosin VI (1:200, Atlas-Sigma HPA035483), Rabbit anti-Histone H3 (tri methyl K9) (1:500, Abcam ab8898), Anti-phospho-Histone H2A.X (Ser139) Antibody produced in mouse (1:500 Sigma-Aldrich, 05-636), Anti-Histone H3 (tri methyl K27) (1:500, Abcam ab6002), Anti-Histone H4 (mono methyl K20) (1:500, Abcam ab177188), Donkey anti-mouse Alexa Fluor 488-conjugated (1:500, Abcam Ab181289), Donkey anti-rabbit Alexa Fluor 647-conjugated (1:500, Abcam Ab181347) and Donkey anti-rabbit Alexa Fluor 488-conjugated antibody (1:500, Abcam Ab181346). Coverslips were mounted on microscope slides with Mowiol (10% (w/v) Mowiol 4-88, 25% (w/v) glycerol, 0.2 M Tris-HCl, pH 8.5), supplemented with 2.5% (w/v) of the anti-fading reagent DABCO (Sigma).

### Confocal Imaging

Cells were visualised using the ZEISS LSM 980 confocal microscope equipped with a Plan-Apochromat 63×/1.4 NA oil immersion objective lens (Carl Zeiss, Cat. No. 420782-9900-000). Three excitation laser lines, 405 nm, 488 nm and 642 nm, were employed to excite the respective fluorophores. Built-in main beam splitters (Carl Zeiss, MBS-405, MBS-488, and MBS-642) directed the laser beams onto the samples.

Emission signals were collected using spectral detection windows of 410–524 nm (Hoechst), 493– 578 nm (green channel), and 564–697 nm (red channel). Detection was achieved using two multi-anode photomultiplier tubes (MA-PMTs) and one gallium arsenide phosphide (GaAsP) detector. The green fluorescence channel was acquired with the GaAsP detector, while blue (Hoechst) and red signals were collected using MA-PMTs. Image acquisition and rendering were performed using ZEN software (Carl Zeiss, ZEN 2.3). All images were then analysed by ImageJ to measure relative intensity in the cytoplasm and nucleus; foci counts and overall fluorescence intensity.

### Widefield Imaging

Widefield images were acquired using a Zeiss Cell Discoverer 7 with a 20x 0.7 NA Plan-Apochromat air objective. LED illumination (385nm and 469nm) corresponding to Hoechst and Alexa-Fluor488 were used with multi bandpass filter sets. Images were captured with a Hamamatsu Orca Flash 4.0 V3 camera using the Zeiss ZEN Blue software (v2.3). All images were then analysed by ImageJ to measure relative intensity in the cytoplasm and nucleus; foci counts and overall fluorescence intensity.

### STORM Imaging

STORM imaging followed our established protocols (41). Cells were seeded on pre-cleaned No 1.5, 25-mm round glass coverslips, placed in 6-well cell culture dishes. Glass coverslips were cleaned by incubating them for 3 hours, in etch solution, made of 5:1:1 ratio of H_2_O: H_2_O_2_ (50 wt. % in H_2_O, stabilized, Fisher Scientific): NH_4_OH (ACS reagent, 28-30% NH_3_ basis, Sigma), placed in a 70°C water bath. Cleaned coverslips were repeatedly washed in filtered water and then ethanol, dried and used for cell seeding. Transfected or non-transfected cells were fixed in pre-warmed 4% (w/v) PFA in PBS and residual PFA was quenched for 15 min with 50 mM ammonium chloride in PBS. Immunofluorescence was performed in filtered sterilised PBS. Cells were permeabilized and simultaneously blocked for 30 min with 3% (w/v) BSA in PBS or TBS, supplemented with 0.1 % (v/v) Triton X-100. Permeabilized cells were incubated for 1h with the primary antibody and subsequently the appropriate fluorophore-conjugated secondary antibody, at the desired dilution in 3% (w/v) BSA, 0.1% (v/v) Triton X-100 in PBS or TBS. The antibody dilutions used were the same as for the normal immunofluorescence protocol (see above), except from the secondary antibodies which were used at 1:250 dilution. Following incubation with both primary and secondary antibodies, cells were washed 3 times, for 10 min per wash, with 0.2% (w/v) BSA, 0.05% (v/v) Triton X-100 in PBS or TBS. Cells were further washed in PBS and fixed for a second time with pre-warmed 4% (w/v) PFA in PBS for 10 min. Cells were washed in PBS and stored at 4 °C, in the dark, in 0.02% NaN3 in PBS, before proceeding to STORM imaging.

Before imaging, coverslips were assembled into the Attofluor® cell chambers (Invitrogen). Imaging was performed in freshly made STORM buffer consisting of 10 % (w/v) glucose, 10 mM NaCl, 50 mM Tris - pH 8.0, supplemented with 0.1 % (v/v) 2-mercaptoethanol and 0.1 % (v/v) pre-made GLOX solution which was stored at 4 ^0^C for up to a week (5.6 % (w/v) glucose oxidase and 3.4 mg/ml catalase in 50 mM NaCl, 10 mM Tris - pH 8.0). All chemicals were purchased from Sigma.

Imaging was undertaken using the Zeiss Elyra PS.1 system. Illumination was from a HR Diode 642 nm (150 mW) and HR Diode 488 nm (100 mW) lasers where power density on the sample was 7-14 kW/cm^2^ and 7-12 kW/cm^2^, respectively.

Imaging was performed under highly inclined and laminated optical (HILO) illumination to reduce the background fluorescence with a 100x NA 1.46 oil immersion objective lens (Zeiss alpha Plan-Apochromat) with a BP 420-480/BP495-550/LP 650 filter. The final image was projected on an Andor iXon EMCCD camera with 25 msec exposure for 20000 frames.

Image processing was performed using the Zeiss Zen software. Where required, two channel images were aligned following a calibration using a calibration using pre-mounted MultiSpec bead sample (Carl Zeiss, 2076-515). For calibration, a 2 μm Z-stack was acquired at 100 nm steps. The channel alignment was then performed in the Zeiss Zen software using the Affine method to account for lateral, tilting and stretching between the channels. The calibration was performed during each day of measurements.

The images were then processed through our STORM analysis pipeline using the Zen software. Single molecule detection and localisation was performed using a 9-pixel mask with a signal to noise ratio of 6 in the “Peak finder” settings while applying the “Account for overlap” function. This function allows multi-object fitting to localise molecules within a dense environment. Molecules were then localised by fitting to a 2D Gaussian.

The render was then subjected to model-based cross-correlation drift correction and detection grouping to remove detections within multiple frames. Typical localisation precision was 20 nm for Alexa-Fluor 647 and 30 nm for Alexa-Fluor 488. The final render was then generated at 10 nm/pixel and displayed in Gauss mode where each localisation is presented as a 2D gaussian with a standard deviation based on its precision. The localisation table was exported as a csv for import into Clus-DoC.

### Clus-DoC

The single molecule positions were exported from Zeiss Zen Black version and imported into the Clus-DoC analysis software (28) (https://github.com/PRNicovich/ClusDoC). The region of interest was determined by the nuclear staining. First the Ripley K function was completed on each channel identifying the r max. The r max was then assigned for DBSCAN if one channel was being analysed or Clus-Doc if two channel colcalisation was being analysed. The MinPts was 5 and a cluster required 10 locations, with smoothing set at 7 nm and epsilon set at the mean localization precision for the dye. All other analyses parameters and colocalization thresholds remained at default settings (28). Data concerning each cluster was exported and plotted using Graphpad.

### Multi-focal Imaging and Particle Tracking Analysis

Cells stably or transiently expressing Halo-tag constructs were labelled for 15 min with 10 nM HaloTag-JF549 ligand, in cell culture medium at 37°C, 5% CO_2_. Cells were washed for 3 times with warm cell culture medium and then incubated for further 30 min at 37°C, 5% CO_2_. Cells were then washed three times in pre-warmed FluoroBrite DMEM imaging medium (ThermoFisher Scientific), before proceeding to imaging.

Single molecule imaging was performed using an aberration-corrected multifocal microscope (acMFM), as described by Abrahamsson et al. (33). Briefly, samples were imaged using 561nm laser excitation, with typical irradiance of 4-6 kW/cm^2^ at the back aperture of a Nikon 100x 1.4 NA objective. Images were relayed through a custom optical system appended to the detection path of a Nikon Ti microscope with focus stabilization. The acMFM detection path includes a diffractive multifocal grating in a conjugate pupil plane, a chromatic correction grating to reverse the effects of spectral dispersion, and a nine-faceted prism, followed by a final imaging lens.

The acMFM produces nine simultaneous, separated images, each representing successive focal planes in the sample, with ca. 20 µm field of view and nominal axial separation of ca. 400nm between them. The nine-image array is digitized via an electron multiplying charge coupled device (EMCCD) camera (iXon Du897, Andor) at up to 32ms temporal resolution, with typical durations of 30 seconds.

3D+t images of single molecules were reconstructed via a calibration procedure, implemented in Matlab (MathWorks), that calculates and accounts for (1) the inter-plane spacing, (2) affine transformation to correctly align each focal plane in the xy plane with respect to each other, and (3) slight variations in detection efficiency in each plane, typically less than ±5-15% from the mean.

Reconstructed data were then subject to pre-processing, including background subtraction, mild deconvolution (3-5 Richardson-Lucy iterations), and/or Gaussian de-noising prior to 3D particle tracking using the MOSAIC software suite (42). Parameters were set where maximum particle displacement was 400 nm and a minimum of 10 frames was required. Tracks were reconstructed, and diffusion constants were extracted via MSD analysis (43) using custom Matlab software assuming an anomalous diffusion model.

### Immunoblot Analysis

The total protein concentration was determined by Bradford Assay (Sigma) following the manufacturer’s instructions. Cell lysates were heat-denatured and resolved by SDS-PAGE. The membrane was probed against the endogenous proteins by incubation with primary Rabbit anti-myosin VI (1:500, Atlas-Sigma HPA035483-100UL), rabbit anti-beta-actin (1:5000, Abcam ab8227), and subsequently secondary Goat anti-rabbit antibody (1:15000 Abcam ab6721) or Goat anti-mouse antibody (1:15000, Abcam ab97023) coupled to horseradish peroxidase. The bands were visualised using the ECL Western Blotting Detection Reagents (Invitrogen) and the images were taken using Syngene GBox system. Images were processed in ImageJ.

### Single molecule FRET

Human Ku70/80 (TP710418) and XRCC4 (TP312684M) proteins were purchased from Origene. Filamentous Actin was purchased from Cytoskeleton Inc (AKF99). DNA oligonucleotides were purchased from IDT each containing a single label of either Atto550 or Alexafluor647. The DNA contained a loop so that only one free end is available for interaction in the assay.

Oligonucleotide 1: AF647- GACCTAGTCGATCGTACGATGCTAC- TTTTAATTTT- GTAGCATCGTACGATCGACTAGGTC

Oligonucleotide 2: Atto550- CGGAGTAACTGCTCTGTTATTTAGC- TAATATATTA- GCTAAATAACAGAGCAGTTACTCCG

Prior to measurements, the DNA was heated to 95°C and then allowed to cool to room temperature to ensure correct annealing. 100 nM DNA was incubated for 15 min at room temperature with 500 nM protein. The sample was diluted to 20 pM in buffer (50 mM Tris.HCl (pH7.5), 50 mM NaCl, 5 mM MgCl_2_, 1 mM DTT) containing the same protein concentration.

Samples were measured in 50 µL on a Precision coverglass #1.5H (Thorlabs). All data was acquired using the EI-Flex Pro HT (Exciting Instruments), a confocal fluorescence spectrometer. All measurements lasted 30 min per well using 520 nm and a 638 nm lasers at 250 µW and 50 µW, respectively, focused in solution (20 µm above the coverslip surface). Data analysis was performed using PhotonFit (Exciting Instruments) with a Burst threshold of 15.

### RNA-seq and analysis

Total RNA was extracted from three replicates of WT, MVI KD, MVI KD+Cisplatin, Scrambled and Scrambled+Cisplatin. Ice cold TRIzol reagent was added to each culture and homogenised. The mixture was then incubated for 5 mins at room temperature then chloroform was added to the lysis and incubated for 3 mins. The samples were then centrifuged at 12,000 x g at 4°C. The colourless aqueous phase was collected. The RNA was then precipitated with incubation for 10 mins with isopropanol before centrifugation for a further 10 mins at 12,000 x g at 4 °C. The pellet was washed in 75% (v/v) ethanol, vortexed and centrifuged for 5 mins at 7500 x g at 4°C. The RNA pellet is air dried for 10 mins. The pellet is then resuspended in 50µL of RNase-free water containing 0.1mM EDTA and incubated at 55°C for 15 mins to allow the RNA to dissolve. The RNA was then quantified using then 260nm absorbance, ensuring the A260/A280 ratio was approximately 2, therefore implying the sample is pure. The sample was then further purified using the RNeasy kit (Qiagen) where the manufacturers protocol was followed exactly. Once the purity and stability had been measured the RNA was then stored at -80°C.

The RNA-seq libraries were prepared with TruSeq RNA Library Prep kit v2 as per protocol instructions. Resulting libraries concentration, size distribution and quality were assessed on a Qubit fluorometer with a dsDNA high sensitivity kit and on an Agilent 2100 bioanalyzer using a DNA 7500 kit. Then libraries were normalized, pooled and quantified with a KAPA Library quantification kit for Illumina platforms on a ABI StepOnePlus qPCR machine, then loaded on a high output flow cell and paired-end sequenced (2×75 bp) on an Illumina NextSeq 550 next generation sequencer (performed at the NYUAD Sequencing Center). The raw FASTQ reads were quality trimmed using Trimmomatic (version 0.36) (44) to trim low-quality bases, systematic base calling errors, as well sequencing adapter contamination. FastQC (www.bioinformatics.babraham.ac.uk/projects/fastqc) was used to assess the quality of the sequenced reads pre/post quality trimming. Only the reads that passed quality trimming in pairs were retained for downstream analysis. The quality trimmed RNAseq reads were aligned to the Homo sapiens GRch38.p4 genome using HISAT2 (version 2.0.4) (45). The resulting SAM alignment files were then converted to BAM format and sorted by coordinate using SAMtools (version 0.1.19) (46). The BAM alignment files were processed using HTseq-count (47) using the reference annotation file to produce raw counts for each sample. The raw counts were then analyzed using the online analysis portal NASQAR (http://nasqar.abudhabi.nyu.edu/) in order to merge, normalize and identify differentially expressed genes by using the START app (48). Differentially expressed genes by at least 2-fold log2(FC)≥1 and adjusted p-value of <0.05 for upregulated genes and log2(FC)≤-1 and adjusted p-value of <0.05 for downregulated genes) between the samples which were then subjected to Gene Ontology (GO) enrichment using ShinyGo v0.60 (http://bioinformatics.sdstate.edu/go/) (49). RNA-Seq data were deposited in the Gene Expression Omnibus (GEO) database under the accession number **TBC**.

### Alkaline Comet Assay

To detect both single, double-stranded breaks and alkali labile sites in DNA, a Comet assay kit (ab238544, Abcam, Cambridge, UK) was used following an adapted version of the manufacturer’s protocol. HeLa cells were washed in ice cold PBS and resuspended to 1×10^6^ cells/mL. The cells were mixed at a 1:10 *v/*v with 37°C comet agarose and dispensed onto the comet slides. The slides were immersed in lysis buffer at 4°C for 90 min followed by unwinding alkaline solution at 4°C for 90 min. The slides underwent electrophoresis in a horizontal gel tank at 18 V and 300 mA for 30 min. The cells were washed in cold deionised water and fixed in cold 70% ethanol. After fixing the cells, the slides were left to air dry at room temperature and stained with Vista Green DNA dye. The cells were imaged on a ZEISS LSM980 using a 10x/0.45 objective with a WD of 2 mm under an FITC filter. Comet tails were analysed using CometScore 2.0.

### Kinase Assay

Substrate Peptides (WT and mutant) were purchased from GL Biochem(Shanghai):

MVI_405 GTKGTVIKVPLKVEQ

MVI_1025 AELISDEAQADLALR

MVI_1155 FLNNSPQQNPAAQIP

ATM kinase was purchased from Eurofins (14933). The ADP-Glo Kinase assay (Promega V9101) was used for the experiments following the manufacturers protocol. Measurements were performed on BMGLabtech Clariostar plate reader.

### Live/Dead Cell stain assay

The kit was purchased from ThermoFisher (L34969). The manufacturers protocol was followed. Cells were treated with the relevant drugs before performing the protocol. Samples were imaged by widefield microscopy.

### Atomic Force Microscopy (AFM)

HeLa cells were seeded on a WillCo-dish and allowed to adhere for 24 h. AFM measurements were performed with a Bruker Bioscope Resolve mounted on an inverted microscope (Nikon Eclipse Ti2) connected to an ORCA-Flash4.0LT (Hamamatsu) camera. Imaging was performed in filtered PBS at room temperature. Pre-calibrated Silicon Tip – Nitride cantilevers (PFQNM-LC-V2 Bruker) were used with a 70 nm tip radius. PeakForce Tapping/PeakForce Capture with a maximum indentation force of 0.5 nN was used. Indentation curves were fitted within Nanoscope Analysis (Bruker) using a cone-sphere model (50).

### MTT Assay

Cell viability was assessed after drug treatments using the 3-(4,5-dimethylthiazol-2-yl)-2,5-diphenyltetrazolium bromide (MTT) assay. Cells were seeded into 96 well plates and treatments were performed for 24 hrs. 10 µl MTT stock (Biotium BT30006)) was added to the culture medium and cells were incubated at 37°C with 5 % CO2 for 4 hrs. Next, the cells were incubated with 200 μl of dimethyl sulfoxide (DMSO; Sigma) for an additional 5 min, and their viability was measured using a microplate reader at 570 nm. The MTT assays were performed at least three times for each concentration of drug and the percentage of surviving cells relative to the non-treated control was determined.

### Graphics

Data fitting and plotting was performed GraphPad Prism. Cartoons were generated using the BioRender software.

## Data Availability

The data supporting the findings of this study are available from the corresponding author on request.

## REFERENCES

1. Jackson SP, Bartek J. The DNA-damage response in human biology and disease. Nature. 2009;461(7267):1071–8.

2. Jeggo PA, Lobrich M. DNA double-strand breaks: their cellular and clinical impact? Oncogene. 2007;26(56):7717–9.

3. Paull TT, Rogakou EP, Yamazaki V, Kirchgessner CU, Gellert M, Bonner WM. A critical role for histone H2AX in recruitment of repair factors to nuclear foci after DNA damage. Curr Biol. 2000;10(15):886–95.

4. Polo SE, Jackson SP. Dynamics of DNA damage response proteins at DNA breaks: a focus on protein modifications. Genes Dev. 2011;25(5):409–33.

5. Reid DA, Keegan S, Leo-Macias A, Watanabe G, Strande NT, Chang HH, et al. Organization and dynamics of the nonhomologous end-joining machinery during DNA double-strand break repair. Proc Natl Acad Sci U S A. 2015;2015(112):E2575–84.

6. Cook AW, Gough RE, Toseland CP. Nuclear myosins - roles for molecular transporters and anchors. J Cell Sci. 2020;2020(11).

7. Gawor M, Lehka L, Lambert D, Toseland CP. Actin from within - how nuclear myosins and actin regulate nuclear architecture and mechanics. J Cell Sci. 2025;2025(3).

8. Hari-Gupta Y, Fili N, Dos Santos A, Cook AW, Gough RE, Reed HCW, et al. Myosin VI regulates the spatial organisation of mammalian transcription initiation. Nat Commun. 2022;13(1):1346.

9. Fili N, Toseland CP. Unconventional Myosins: How Regulation Meets Function. Int J Mol Sci. 2019;2019(1).

10. Cook AW, Toseland CP. The roles of nuclear myosin in the DNA damage response. J Biochem. 2020.

11. Fili N, Hari-Gupta Y, Dos Santos A, Cook A, Poland S, Ameer-Beg SM, et al. NDP52 activates nuclear myosin VI to enhance RNA polymerase II transcription. Nat Commun. 2017;8(1):1871.

12. Shahid-Fuente IW, Toseland CP. Myosin in chromosome organisation and gene expression. Biochem Soc Trans. 2023;51(3):1023–34.

13. Vreugde S, Ferrai C, Miluzio A, Hauben E, Marchisio PC, Crippa MP, et al. Nuclear myosin VI enhances RNA polymerase II-dependent transcription. Mol Cell. 2006;23(5):749–55.

14. Hari-Gupta Y, Lambert D, Shahid-Fuente IW, Fili N, Santos Ád, Sprules A, et al. Myosin VI orchestrates estrogen-driven gene expression in breast cancer cells. bioRxiv. 2025:2025.11.13.688007.

15. Cho SJ, Chen X. Myosin VI is differentially regulated by DNA damage in p53- and cell type-dependent manners. J Biol Chem. 2010;285(35):27159–66.

16. Jung EJ, Liu G, Zhou W, Chen X. Myosin VI is a mediator of the p53-dependent cell survival pathway. Mol Cell Biol. 2006;26(6):2175–86.

17. Shi J, Hauschulte K, Mikicic I, Maharjan S, Arz V, Strauch T, et al. Nuclear myosin VI maintains replication fork stability. Nat Commun. 2023;14(1):3787.

18. Chai B, Huang J, Cairns BR, Laurent BC. Distinct roles for the RSC and Swi/Snf ATP-dependent chromatin remodelers in DNA double-strand break repair. Genes Dev. 2005;19(14):1656–61.

19. Dos Santos A, Cook AW, Gough RE, Schilling M, Olszok NA, Brown I, et al. DNA damage alters nuclear mechanics through chromatin reorganization. Nucleic Acids Res. 2020.

20. Dos Santos A, Toseland CP. Regulation of Nuclear Mechanics and the Impact on DNA Damage. Int J Mol Sci. 2021;2021(6).

21. Gonzalez-Novo R, Zamora-Carreras H, Armesto M, de Lope-Planelles A, Lopez-Menendez H, Roda-Navarro P, et al. 3D environment favors persistent changes in cell functions and altered morphology, wrinkling, and biomechanical signature of the nucleus. Cell Rep Phys Sci. 2026;7(2):103116.

22. Ziv Y, Bielopolski D, Galanty Y, Lukas C, Taya Y, Schultz DC, et al. Chromatin relaxation in response to DNA double-strand breaks is modulated by a novel ATM- and KAP-1 dependent pathway. Nat Cell Biol. 2006;8(8):870–6.

23. Dunn TA, Chen S, Faith DA, Hicks JL, Platz EA, Chen Y, et al. A novel role of myosin VI in human prostate cancer. Am J Pathol. 2006;169(5):1843–54.

24. Puri C, Chibalina MV, Arden SD, Kruppa AJ, Kendrick-Jones J, Buss F. Overexpression of myosin VI in prostate cancer cells enhances PSA and VEGF secretion, but has no effect on endocytosis. Oncogene. 2010;29(2):188–200.

25. Wang D, Zhu L, Liao M, Zeng T, Zhuo W, Yang S, et al. MYO6 knockdown inhibits the growth and induces the apoptosis of prostate cancer cells by decreasing the phosphorylation of ERK1/2 and PRAS40. Oncol Rep. 2016;36(3):1285–92.

26. Wang H, Wang B, Zhu W, Yang Z. Lentivirus-Mediated Knockdown of Myosin VI Inhibits Cell Proliferation of Breast Cancer Cell. Cancer Biother Radiopharm. 2015;30(8):330–5.

27. Yoshida H, Cheng W, Hung J, Montell D, Geisbrecht E, Rosen D, et al. Lessons from border cell migration in the Drosophila ovary: A role for myosin VI in dissemination of human ovarian cancer. Proc Natl Acad Sci U S A. 2004;101(21):8144–9.

28. Pageon SV, Nicovich PR, Mollazade M, Tabarin T, Gaus K. Clus-DoC: a combined cluster detection and colocalization analysis for single-molecule localization microscopy data. Mol Biol Cell. 2016;27(22):3627–36.

29. Malkusch S, Endesfelder U, Mondry J, Gelleri M, Verveer PJ, Heilemann M. Coordinate-based colocalization analysis of single-molecule localization microscopy data. Histochem Cell Biol. 2012;137(1):1–10.

30. Banath JP, Klokov D, MacPhail SH, Banuelos CA, Olive PL. Residual gammaH2AX foci as an indication of lethal DNA lesions. BMC Cancer. 2010;10:4.

31. Martin M, Terradas M, Hernandez L, Genesca A. gammaH2AX foci on apparently intact mitotic chromosomes: not signatures of misrejoining events but signals of unresolved DNA damage. Cell Cycle. 2014;13(19):3026–36.

32. Schmid TE, Zlobinskaya O, Multhoff G. Differences in Phosphorylated Histone H2AX Foci Formation and Removal of Cells Exposed to Low and High Linear Energy Transfer Radiation. Curr Genomics. 2012;13(6):418–25.

33. Abrahamsson S, Chen J, Hajj B, Stallinga S, Katsov AY, Wisniewski J, et al. Fast multicolor 3D imaging using aberration-corrected multifocus microscopy. Nat Methods. 2013;10(1):60–3.

34. Dos Santos A, Fili N, Hari-Gupta Y, Gough RE, Wang L, Martin-Fernandez M, et al. Binding partners regulate unfolding of myosin VI to activate the molecular motor. Biochem J. 2022;479(13):1409–28.

35. Bastianello G, Porcella G, Beznoussenko GV, Kidiyoor G, Ascione F, Li Q, et al. Cell stretching activates an ATM mechano-transduction pathway that remodels cytoskeleton and chromatin. Cell Rep. 2023;42(12):113555.

36. Cook A, Hari-Gupta Y, Toseland CP. Application of the SSB biosensor to study in vitro transcription. Biochem Biophys Res Commun. 2018;496(3):820–5.

37. Fili N, Hari-Gupta Y, Aston B, Dos Santos A, Gough RE, Alamad B, et al. Competition between two high- and low-affinity protein-binding sites in myosin VI controls its cellular function. J Biol Chem. 2020;295(2):337–47.

38. Kim JA, Kruhlak M, Dotiwala F, Nussenzweig A, Haber JE. Heterochromatin is refractory to gamma-H2AX modification in yeast and mammals. J Cell Biol. 2007;178(2):209–18.

39. Wollscheid HP, Biancospino M, He F, Magistrati E, Molteni E, Lupia M, et al. Diverse functions of myosin VI elucidated by an isoform-specific alpha-helix domain. Nat Struct Mol Biol. 2016;23(4):300–8.

40. Große-Berkenbusch A, Hettich J, Kuhn T, Fili N, Cook AW, Hari-Gupta Y, et al. Myosin VI moves on nuclear actin filaments and supports long-range chromatin rearrangements. bioRxiv. 2020:2020.04.03.023614.

41. Dos Santos A, Gough RE, Wang L, Toseland CP. Measuring Nuclear Organization of Proteins with STORM Imaging and Cluster Analysis. Methods Mol Biol. 2022;2476:293–309.

42. Sbalzarini IF, Koumoutsakos P. Feature point tracking and trajectory analysis for video imaging in cell biology. J Struct Biol. 2005;151(2):182–95.

43. Aaron J, Wait E, DeSantis M, Chew TL. Practical Considerations in Particle and Object Tracking and Analysis. Curr Protoc Cell Biol. 2019;83(1):e88.

44. Bolger AM, Lohse M, Usadel B. Trimmomatic: a flexible trimmer for Illumina sequence data. Bioinformatics. 2014;30(15):2114–20.

45. Kim D, Langmead B, Salzberg SL. HISAT: a fast spliced aligner with low memory requirements. Nat Methods. 2015;12(4):357–60.

46. Li H, Handsaker B, Wysoker A, Fennell T, Ruan J, Homer N, et al. The Sequence Alignment/Map format and SAMtools. Bioinformatics. 2009;25(16):2078–9.

47. Anders S, Pyl PT, Huber W. HTSeq--a Python framework to work with high-throughput sequencing data. Bioinformatics. 2015;31(2):166–9.

48. Nelson JW, Sklenar J, Barnes AP, Minnier J. The START App: a web-based RNAseq analysis and visualization resource. Bioinformatics. 2017;33(3):447–9.

49. Ge SX, Jung D, Yao R. ShinyGO: a graphical enrichment tool for animals and plants. Bioinformatics. 2019.

50. Bertolio R, Napoletano F, Del Sal G. Dynamic links between mechanical forces and metabolism shape the tumor milieu. Curr Opin Cell Biol. 2023;84:102218.

