## Supplementary Data for "Nuclear Myosin VI stabilises Ku-associated DNA ends during nonhomologous end joining"

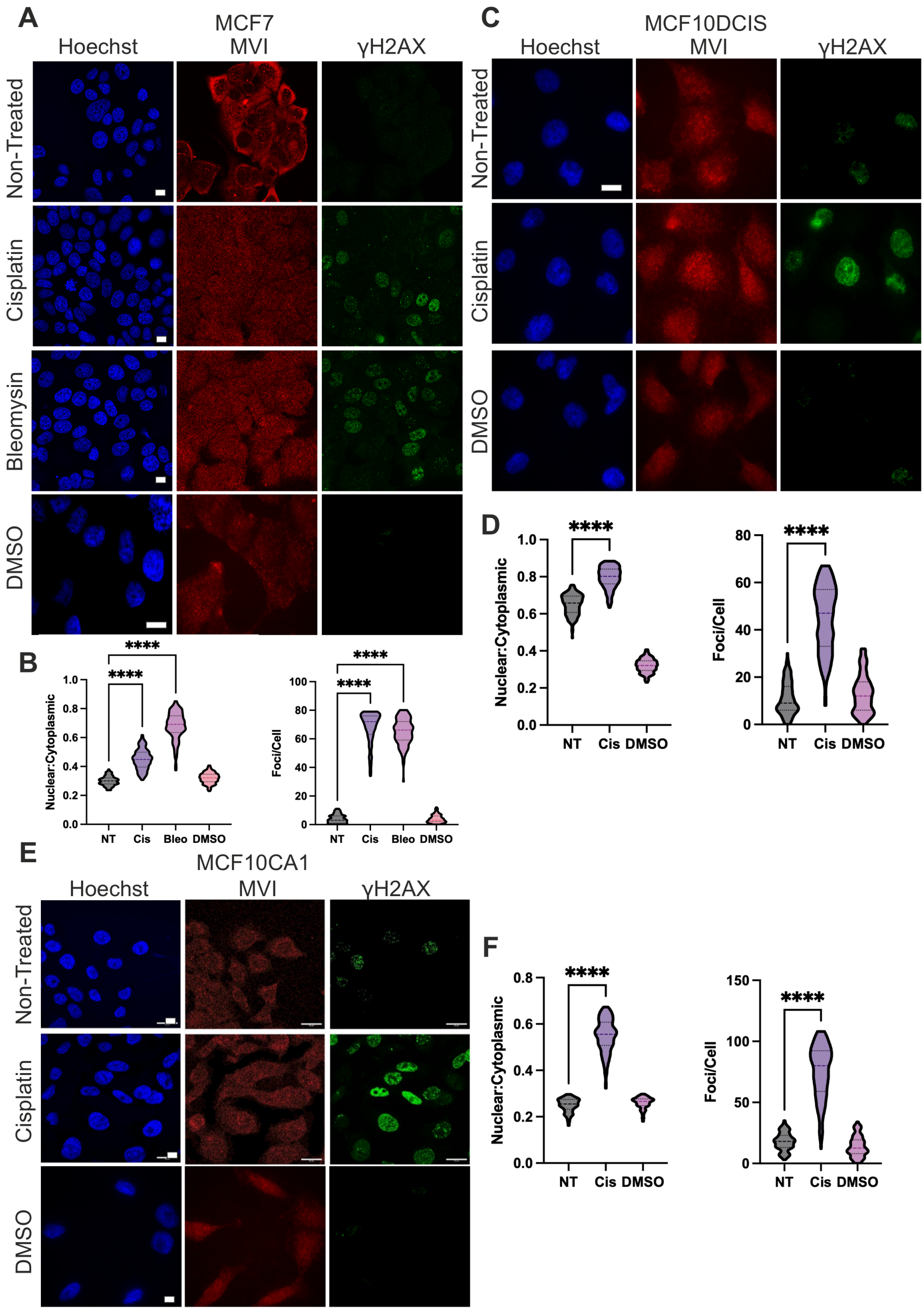


**Supplementary Figure 1: The nuclear accumulation of myosin VI following DNA damage in breast cancer model cell lines.**
Representative Immunofluorescence staining against MVI (red), γH2AX (green) and DNA (blue), and Quantification of nuclear MVI and number of γH2AX foci for: MCF7 cells (A) and (B), MCF10DCIS (C) and (D), and MCF10CA1 cells (E) and (F). Where stated, cells were treated with cisplatin (65 μM 4 h) and bleomycin (0.5 μM 4 hr) treated cells (Scale bar 10 μm). 150 cells per condition from three independent experiments. ****p <0.0001 by one-way ANOVA and Tukey's multiple comparisons test.

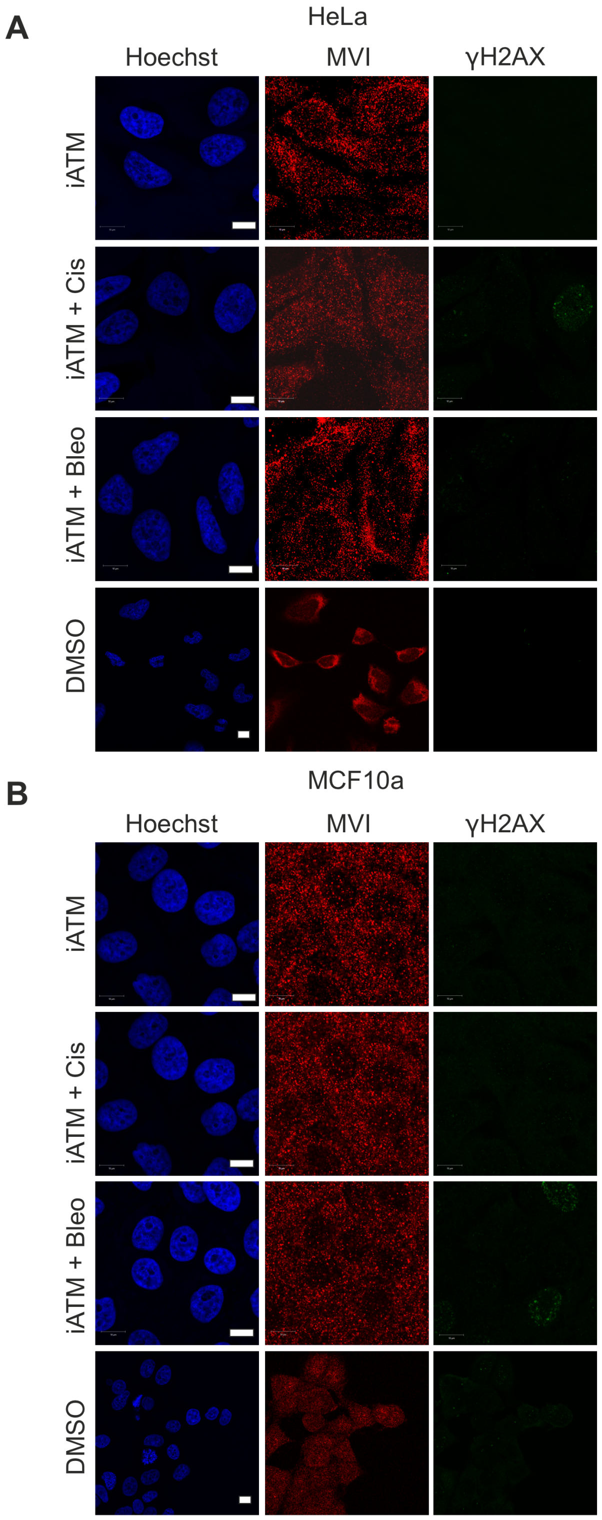


**Supplementary Figure 2: The nuclear accumulation of myosin VI following DNA damage with ATM kinase inhibitor.**
Representative Immunofluorescence staining against MVI (red), γH2AX (green) and DNA (blue) in (A) HeLa cells and (B) MCF10a cells for ATM kinase inhibitor (iATM), (iATM+cisplatin (65 μM 4 h) and iATM+bleomycin (0.5 μM 4 hr) treated cells (Scale bar 10 μm).

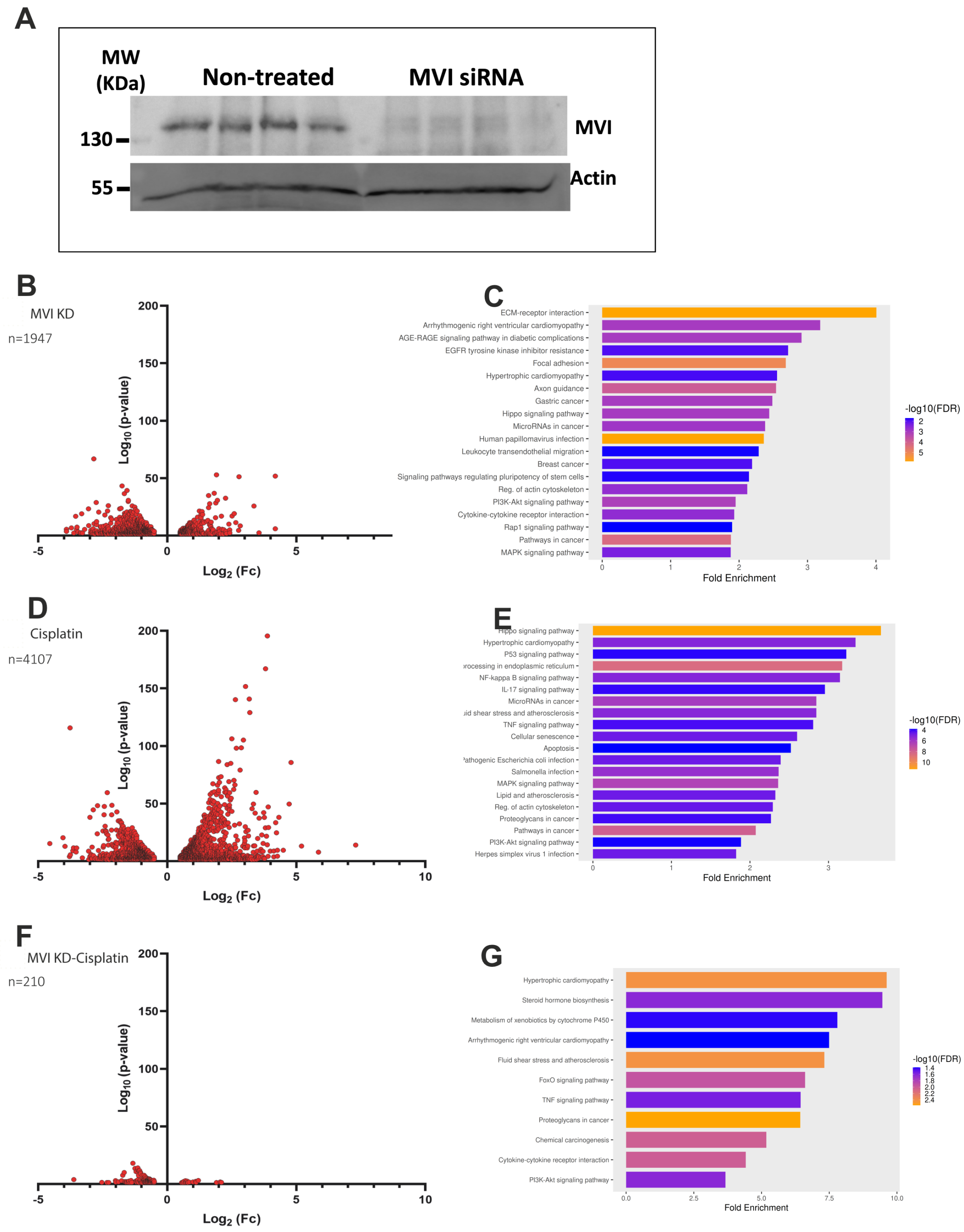


**Supplementary Figure 3: DNA damage induced gene expression changes**

(A) Western blot for the non-treated and MVI siRNA knockdown cells. Uncropped images are in Supplementary Figure 4. (B) Volcano plot of differentially expressed genes from RNA-seq following MVI siRNA knockdown in HeLa cells. (C) GO terms with arranged by fold enrichment for pathway terms colour-coded by -log10(FDR) corresponding to the differentially expressed genes following MVI knockdown. (D) Volcano plot of differentially expressed genes from RNA-seq following 25 μM cisplatin treatment for 24h in HeLa cells. (E) GO terms with arranged by fold enrichment for pathway terms colour-coded by -log10(FDR) corresponding to the differentially expressed genes following cisplatin treatment. (F) Volcano plot of differentially expressed genes from RNA-seq following MVI siRNA knockdown and treated with cisplatin for 24 h in HeLa cells. The differential expression analysis compared conditions in B to those occurring following knockdown and then cisplatin-treatment. (G) GO terms with arranged by fold enrichment for pathway terms colour-coded by -log10(FDR) corresponding to the differentially expressed genes following MVI knockdown-Cisplatin treatment.

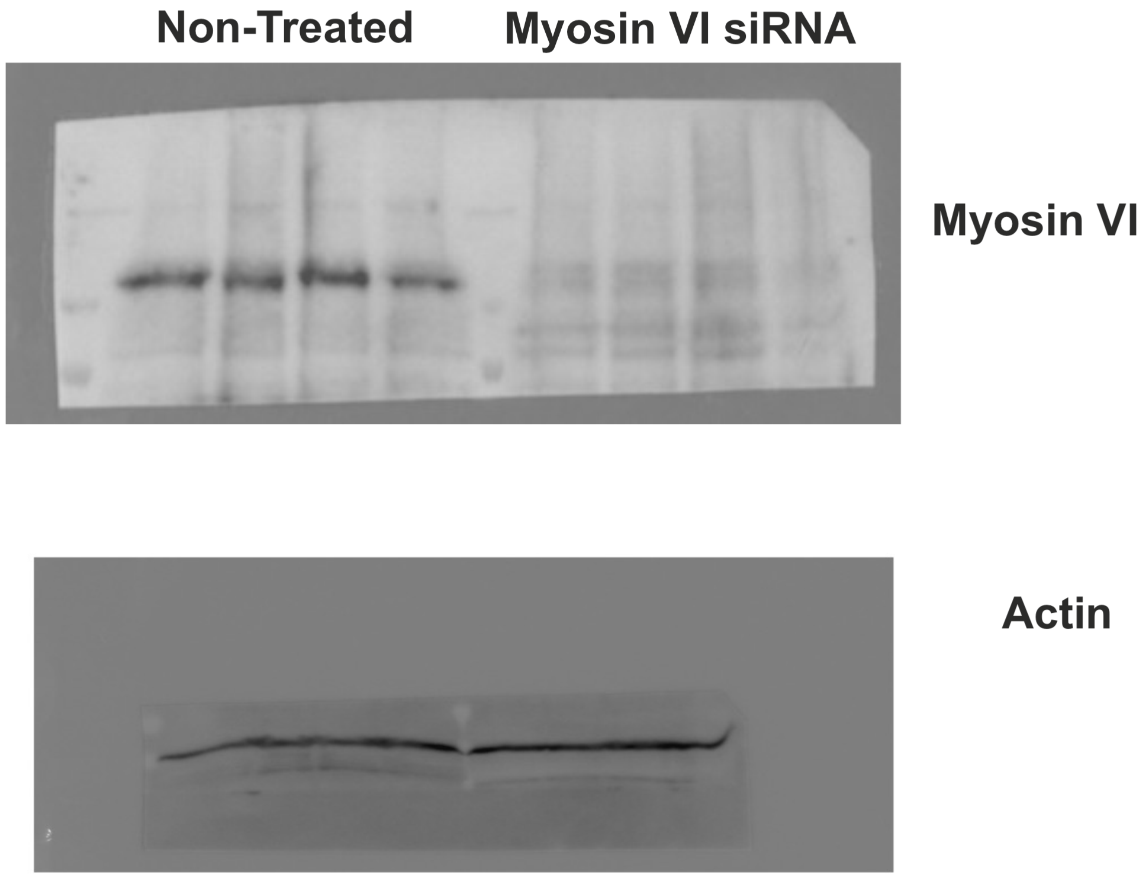


**Supplementary Figure 4: Full view of western-blot.**
Uncropped Western blot images for the non-treated and MVI siRNA knockdown cells in Supplementary Figure 3.

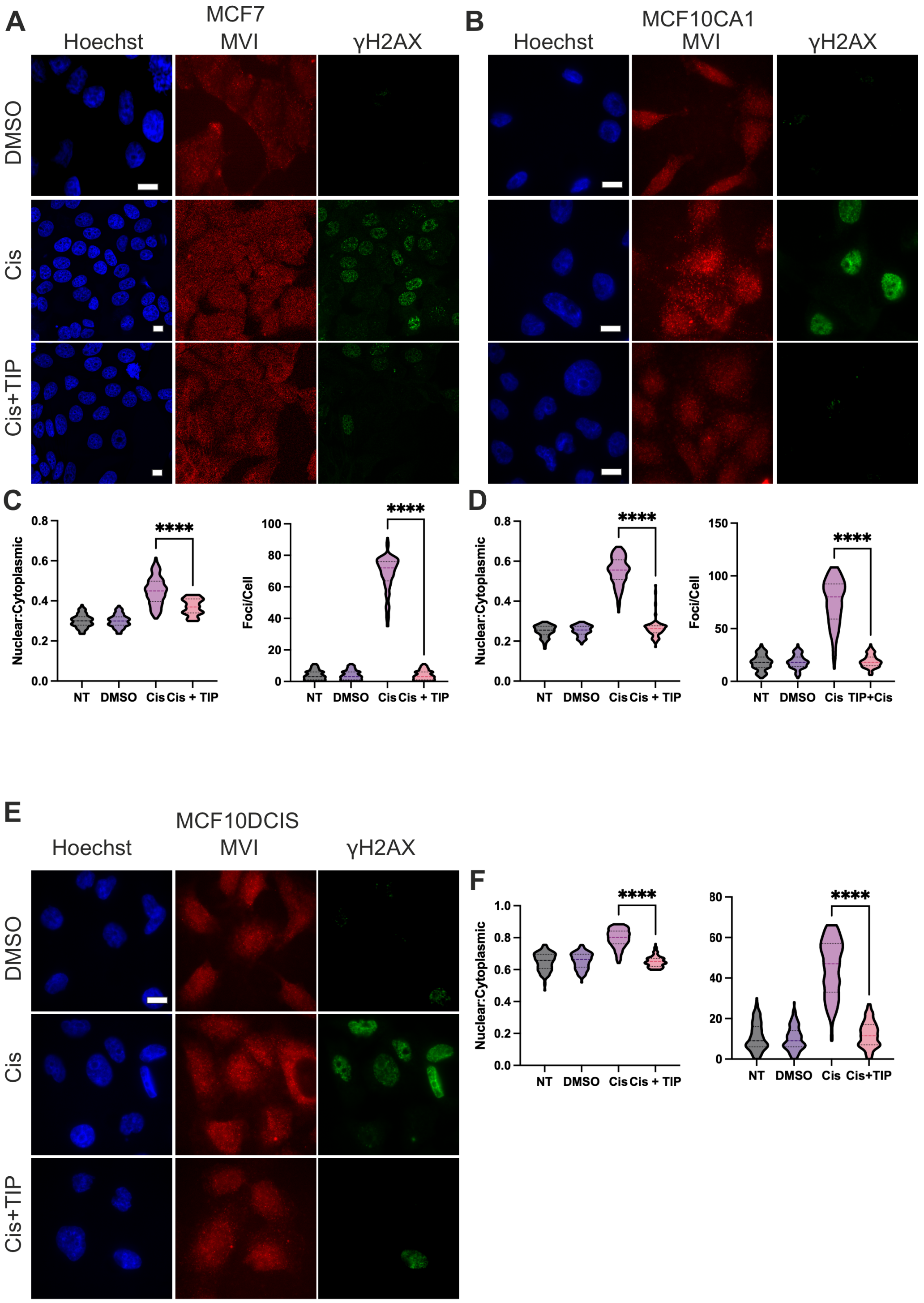


**Supplementary Figure 5: Myosin VI perturbation disrupts γH2AX signalling.** Representative Immunofluorescence staining against MVI (red), γH2AX (green) and DNA (blue) in (A) MCF7 cells and (B) MCF10CA1 cells and (E) for DMSO control, Cisplatin-Treated (65 μM 4 h) and TIP+cisplatin (65 μM 4 h) (Scale bar 10 μm). (C), (D) and (F) Quantification of nuclear MVI and number of γH2AX foci for MCF7, MCF10CA1 and MCF10DCIS cells, respectively. 150 cells per condition from three independent experiments. ****p <0.0001 by one-way ANOVA and Tukey's multiple comparisons test.

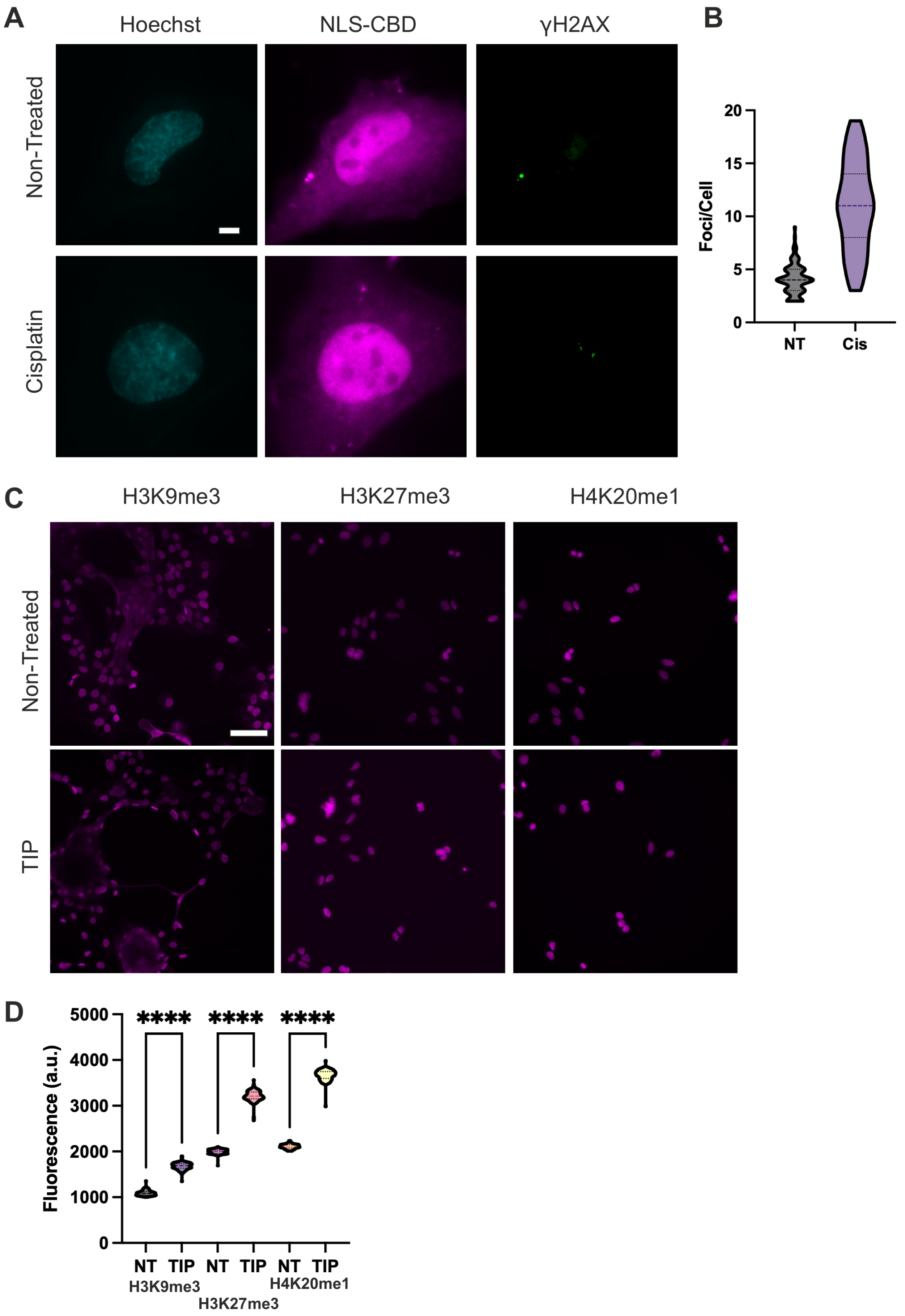


**Supplementary Figure 6: Specific targeting of the nuclear MVI pool and perturbation impact on chromatin**

(A) Representative Immunofluorescence staining against γH2AX (green) and DNA (cyan) in HeLa cells following transfection with Halo-NLS-CBD (magenta). Cells were transfected for 48 h before cisplatin treatment for 4 h at 65 µM. (Scale bar 10 µm). (B) Quantification of γH2AX foci in both conditions.

(C) Representative Immunofluorescence staining against H3K9me3, H3K27me3 and H4K20me1 in HeLa cells for non-treated and TIP treated conditions (25 μM 4 h) (Scale bar 100 μm). (D) Quantification of fluorescence intensity for the experiment in (C). 150 cells per condition from three independent experiments. ****p <0.0001 by one-way ANOVA and Tukey's multiple comparisons test.

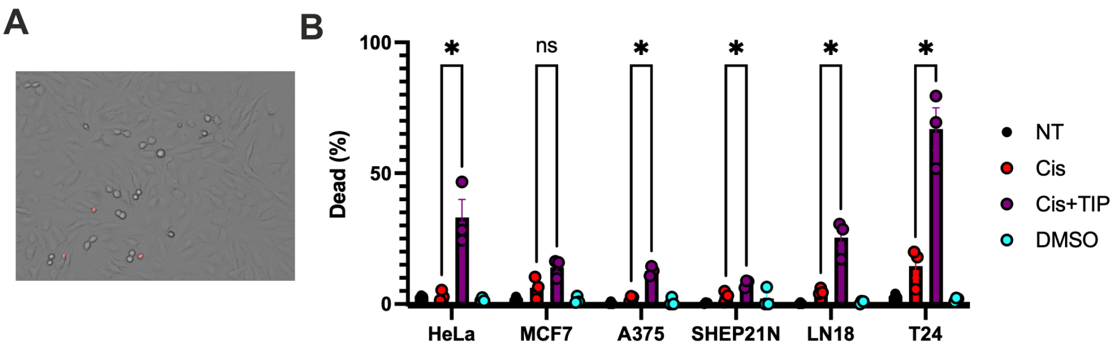


**Supplementary Figure 7: Myosin VI inhibition sensitises cells to DNA damage agents leading to increased cell death.**
(A) Example image identifying dead cells within the field of view. (B) Quantification of the percentage of dead cells for each cell line under the stated conditions. The datapoints represent the mean from three biological repeats consisting of at least 1200 cells. *p <0.05 by one-way ANOVA and Tukey's multiple comparisons test. Error bars represent SEM.
